# Polarity-dependent expression and localization of secretory glucoamylase mRNA in filamentous fungal cells

**DOI:** 10.1101/2023.10.12.562120

**Authors:** Yuki Morita, Kaoru Takegawa, Brett M. Collins, Yujiro Higuchi

## Abstract

In multinuclear and multicellular filamentous fungi little is known about how mRNAs encoding secreted enzymes are transcribed and localized spatiotemporally. To better understand this process we visualized mRNA encoding GlaA, a glucoamylase secreted in large amounts by the industrial filamentous fungus *Aspergillus oryzae*, in living cells by the MS2 system. We found that *glaA* mRNA was significantly transcribed and localized near the hyphal tip and septum, which are the sites of protein secretion. We also revealed that *glaA* mRNA exhibits long-range dynamics in the vicinity of the endoplasmic reticulum (ER) in a manner that is dependent on the microtubule motor proteins kinesin-1 and kinesin-3, but independent of early endosomes. Moreover, we elucidated that the regulatory mechanisms of *glaA* mRNA by stress granules and processing bodies were different under high temperature and ER stress. Collectively, this study uncovers a dynamic regulatory mechanism of mRNA encoding a secretory enzyme in filamentous fungi.

## INTRODUCTION

The mechanism controlling cellular mRNA localization is necessary for the spatiotemporal regulation of gene expression and is a widely conserved biological process from prokaryotes to eukaryotes.^1-5^ This mechanism is thought to efficiently regulate the subsequent function of translated proteins by promoting their local or compartment specific biosynthesis.^6^ The mechanism of mRNA localization has been extensively studied in neurons, and in model organisms such as *Drosophila melanogaster* and *Saccharomyces cerevisiae*, and has been found to be important for asymmetric biological phenomena including cell polarity formation, cell cycle and differentiation.^7^ In multicellular organisms, loss of mRNA localization is known to cause significant developmental and growth abnormalities.^8-10^ The molecular mechanisms of mRNA localization have been intensively analyzed in relation to their importance in a variety of cellular activities, but it is still unclear how mRNA localization contributes to a wide range of biological phenomena.

Certain mRNAs are transported in an untranslated state using molecular motors that travel along the cytoskeleton, but mRNA localization mechanisms differ among species.^11,12^ For example, *ASH1* mRNA in *S. cerevisiae* is transported to budding daughter cells by binding to the myosin-V Myo4, via the She2p-She3p complex.^13-17^ In the dimorphic fungus *Ustilago maydis*, certain mRNAs bind to early endosomes (EEs), which are moved along microtubules by motor proteins kinesin-3 and dynein, via a complex containing Rrm4, Upa1, Upa2, Grp1, Pab1, etc., and are transported in both directions within hyphal cells.^3,18,19^ The mRNAs bound to the EE are then transported in the form of polysomes with multiple ribosomes that are translationally active.^20^ In neurons, mRNAs are known to be transported in association with organelles including late endosomes and lysosomes.^21,22^ Recently, a sophisticated complex called FERRY has been discovered, which binds to EEs via the small GTPase Rab5, and recruits and transports mRNAs including those that encode mitochondrial proteins.^23,24^ However, these specific protein-mediated mechanisms are not well conserved in fungi, suggesting that mRNA localization machinery has diversified through the process of species evolution and differentiation.

mRNAs that are translationally repressed in the intracellular trafficking process may be subject to a variety of controls before translation.^25-27^ In such regulation, ribonucleoprotein (RNP) granules play an important role in mRNA storage, degradation, and translational repression, and well-studied RNP granules are known as processing bodies (P-bodies, PBs) and stress granules (SGs).^28-31^ These are membraneless organelles formed by liquid-liquid phase separation, in which untranslated mRNAs and RNA-binding proteins are incorporated. As PB components, deadenylases (e.g., Ccr4-Not, Lsm1-7 complex), decapping-related enzymes (e.g., Dcp1, Dcp2, Edc3, Edc4, Dhh1) and 5′-to-3′ exoribonuclease (e.g., Xrn1) are known; however, PBs do not contain ribosomes or translation initiation factors, suggesting they contribute primarily to mRNA degradation.^32^ On the other hand, it is known that mRNA is released from PBs and can be incorporated into SGs. This can initiate translation during stress, and thus may lead mRNA to re-initiate translation.^33^ SGs are known to form transiently during various environmental stresses and are composed of untranslated mRNA, ribosomal small subunits, translation initiation factors (e.g., eIF4GI, eIF4GII, eIF4E), poly(A) binding protein (e.g., Pab1/PABP), ubiquitin protease cofactor (e.g., Bre5/Nxt3/G3BP1) and other RNA binding proteins (e.g., Pub1/TIA-1, Pbp1/Ataxin-2) as well as translation initiation of aggregated mRNAs.^31,34,35^ However, it has been reported that the components of the SG differ depending on the type of stress, and budding yeast does not contain translation initiation factors or ribosomal small subunits during glucose starvation.^36^ The response to environmental stresses such as heat, osmotic, oxidative and UV stresses has been well studied, but intracellular stresses such as endoplasmic reticulum (ER) stress and lysosomal damage have also been reported to lead to SG formation.^35,37,38^

Protein secretion is important for cell growth, homeostasis, cell division, adaptation to the external environment and controlling cell-cell interactions across species.^39,40^ Secreted enzymes are generally synthesized in the ER, glycosylated in the Golgi apparatus and transported within constitutive or regulated secretory vesicles to be released at the cell surface. Most secreted enzymes that follow such pathways have their mRNAs recruited to the ER by the signal recognition particle (SRP) pathway.^41,42^ mRNAs for secreted enzymes encode an ER-localizing signal peptide consisting of about 10 amino acid residues. When this sequence is translated in the cytoplasm, SRP binds to the ER-localizing signal peptide and prevents translation elongation by the ribosome. mRNAs encoding secreted enzymes in the translationally repressed state are thus recruited to the ER by binding of SRP and SRP receptors. The secreted enzyme mRNA tethered to the ER membrane forms polysomes to be translated and the peptide chains translocate into the ER lumen via the Sec61 complex, a translocon complex that is adjacent to the SRP receptor. It is thought that secreted enzyme mRNAs may be transported near the ER membrane to maximize translation efficiency of secreted enzymes and membrane proteins,^18^ although in general there is little known about where secreted enzyme mRNAs are localized in the cell.

*Aspergillus oryzae* is a filamentous fungus widely used in the fermentation and brewing industries, and secretes large amounts of diverse and useful hydrolytic enzymes in a safe manner.^43,44^ However, because filamentous fungi are multinucleated and multicellular compared to yeast and other uninucleate single-celled eukaryotes, the details of the intracellular mechanisms from transcription to translation and transport to secretion have not yet been fully elucidated. Among the secreted proteins produced by *A. oryzae*, amylase is particularly well expressed, and its transcription system and secretory pathway have been well studied.^44-47^ The amylase family includes α-amylases (AmyA, AmyB, AmyC), which hydrolyze the α-1,4 bonds of starch randomly, and glucoamylases (GlaA, GlaB), which hydrolyze glucose from the nonreducing end. Transcriptional induction of these enzymes is regulated by a carbon catabolite repression mechanism, with expression suppressed in the presence of glucose and highly expressed in the presence of maltose.^48-52^ Studies of amylase AmyB have shown it to localize to the hyphal tip and septum, and these have been proposed to be the major sites of secretion.^45^ Furthermore, visualization of α-amylase mRNA by single-molecule fluorescence in situ hybridization (smFISH) revealed a prominent mRNA fluorescence signal in the region around 50 µm from the hyphal tip.^53^ In addition to these results, mRNA localization in *A. oryzae* differs according to intracellular protein localization, suggesting that the localization of secreted enzyme mRNA may have an important role in the efficiency of protein secretion.^54^

In this study we have examined the glucoamylase (GlaA) protein, which is secreted in large amounts by *A. oryzae*, and analyzed the localization mechanism of its mRNA. We found that *glaA* mRNA is selectively transcribed in the nucleus near the secretory sites and then localized near the hyphal tip and septum. We show that *glaA* mRNA localizes mainly to the ER, with dynamics that are regulated by kinesin-1 and kinesin-3 Furthermore, the results suggest that secreted enzyme mRNAs are differentially regulated depending on heat or ER stresses.

## RESULTS

### Visualization of *glaA* mRNA by MS2 system

The MS2 system enables visualization of target mRNAs at the single molecule level using an RNA sequence derived from bacteriophage MS2 (MS2-binding site, MBS) and an RNA-binding protein that specifically binds to MBS (MS2-coat protein, MCP), and is a technology widely applied in animal and plant cells.^1,14^ In this study, we generated a strain in which the MS2 system was applied to the glucoamylase-encoding *glaA* gene (*glaA*-MS2 strain) in order to analyze the localization mechanism of mRNA encoding the secreted enzyme in *A. oryzae* living cells. We inserted a 36×MBS sequence downstream of the *glaA* gene and co-expressed the nuclear localization signal (NLS)-fused MCP-2×EGFP to clearly visualize dot-like fluorescence that might indicate *glaA* mRNAs (Figures 1A and 1B).

**Figure 1.**
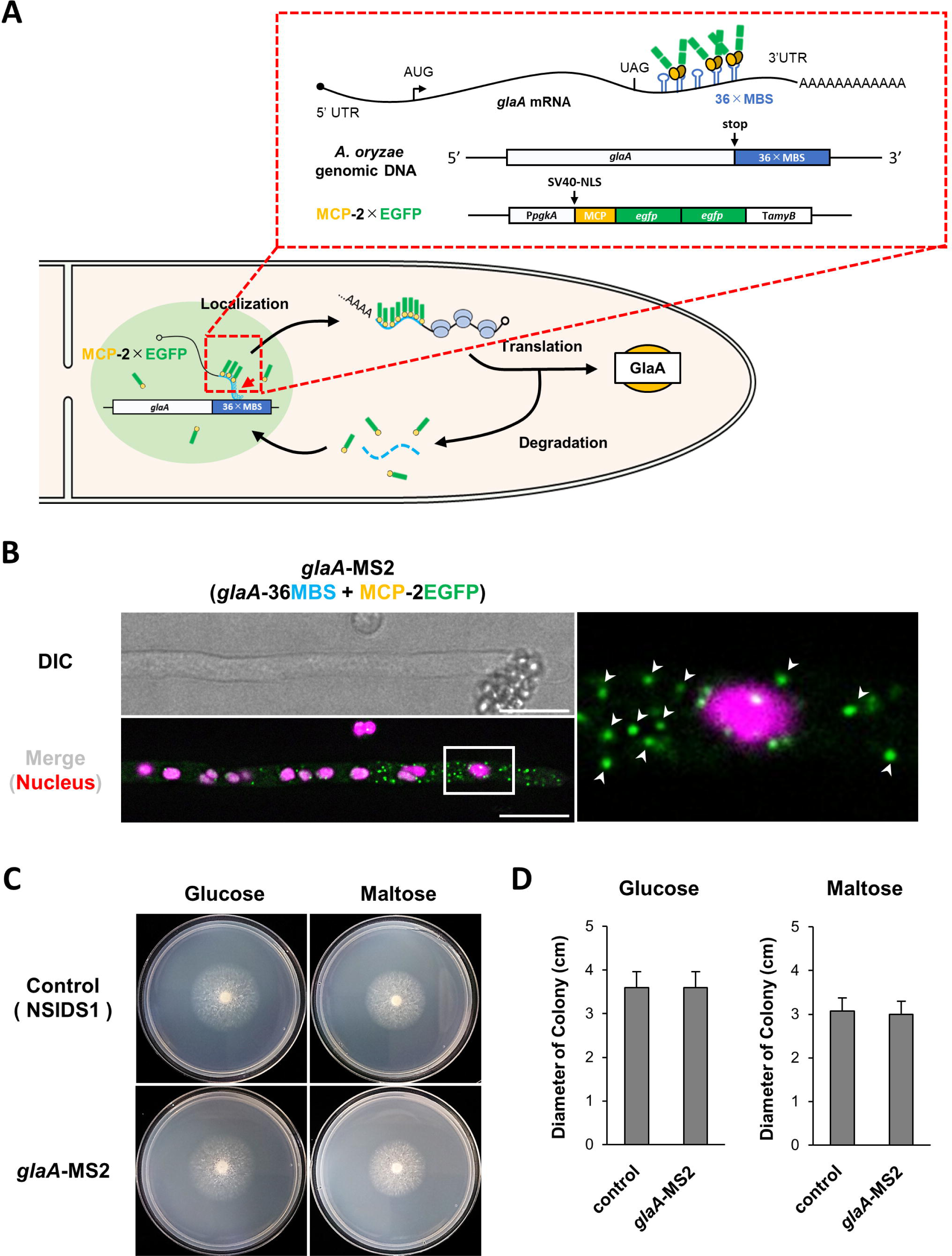
Observation of *glaA* mRNA by using the MS2 system in *A. oryzae*. (A) Schematic diagram of MS2 system for visualizing *glaA* mRNA in *A. oryzae*. (B) Colocalization of *glaA* mRNA (green) and nuclei (magenta) in the apical cell. White arrowheads show *glaA* mRNA foci in the enlarged image on the right. (C) Growth tests of the control (NSlDS1) and *glaA*-MS2 strains. (D) Diameters of the colony in each culture condition. Error bars indicate the standard deviation of the mean (n = 3).

After applying the MS2 system to *A. oryzae glaA*, we analyzed the effects on mycelial growth, gene expression and enzyme activity. Although various carbon sources affect *glaA* gene expression, no significant differences in growth rate or colony morphology were observed between control and *glaA*-MS2 strains under several culture conditions (Figures 1C, 1D and S1A-S1B). The *glaA*-MS2 strain was reduced to about 34% of the control strain with respect to the mRNA level of the *glaA* gene (Figure S1C), and was also reduced to about 39% with respect to secreted glucoamylase activity (Figure S1D and S1E). These results indicate that the introduction of the MS2 system into *glaA* does not affect growth, but reduces the levels of mRNA and enzyme activity. The reduction of intracellular mRNA levels of target genes by the introduction of the MS2 system has been reported previously and has been analyzed to the extent that the localization of target mRNAs is not affected.^55-58^

### Effects of MS2 system for localization and expression of *glaA* mRNA

The dot-like fluorescence visualized by the MS2 system was analyzed by smFISH to see if it actually indicates the subcellular localization of *glaA* mRNA. First, we conducted smFISH using fluorescent probes specific for *glaA* and MBS, and found that *glaA* and MBS mRNA fluorescence co-localized in the cytoplasm (Figure 2A). Furthermore, we confirmed that *glaA* mRNA fluorescence derived from the binding of MCP-2×EGFP to MBS and mRNA fluorescence derived from the *glaA*-specific probe by smFISH showed co-localization (Figure 2B).

**Figure 2.**
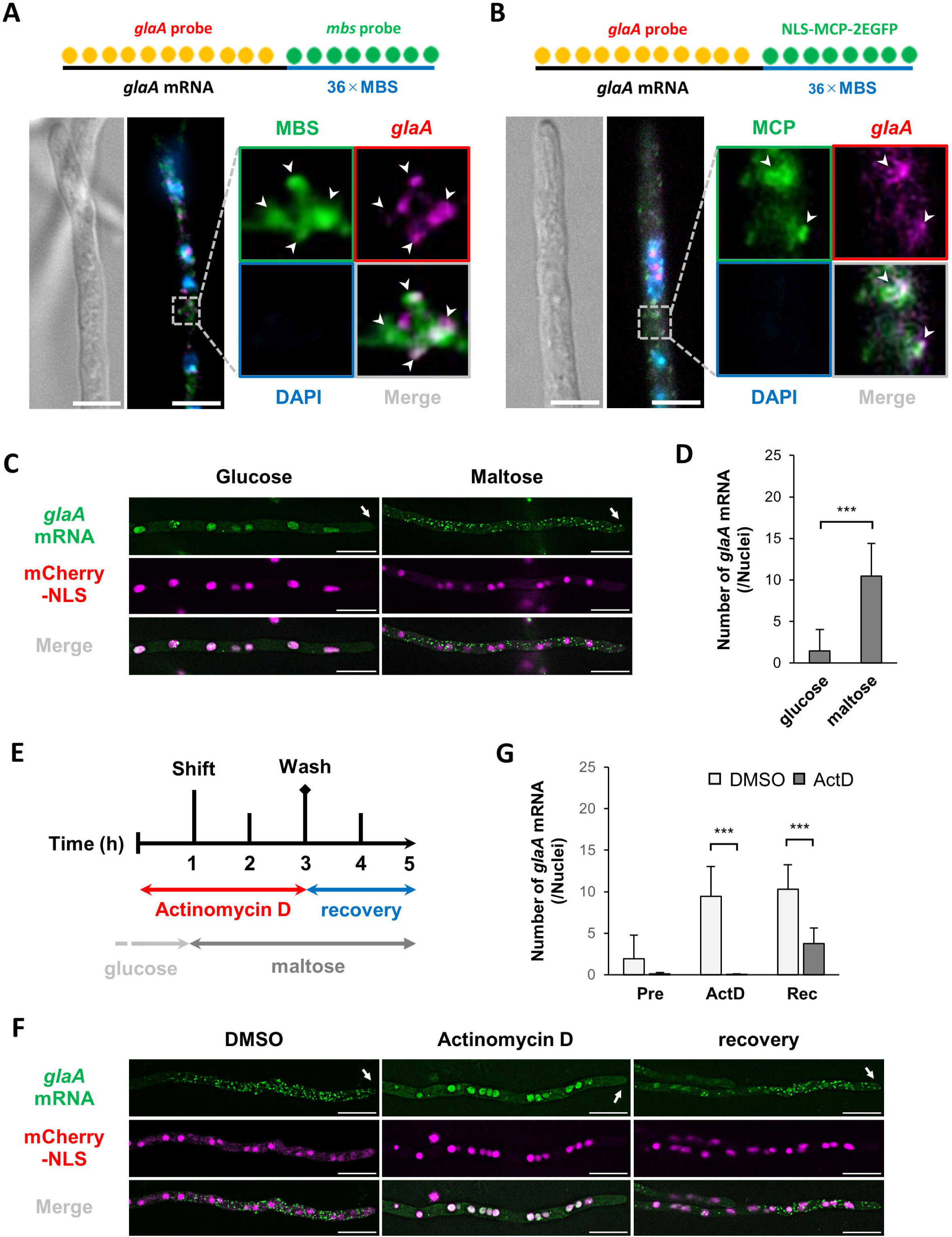
Effects of MS2 system for localization and expression of *glaA* mRNA. (A and B) Colocalization analysis of *glaA* mRNAs, MBS mRNAs and MCP-2×EGFP foci by smFISH. White arrowheads show colocalized foci. Scale bars, 5 µm. (C) Colocalization of *glaA* mRNAs (green) and nuclei (magenta).White arrows indicate the hyphal tip. Scale bars, 10 µm. (D and G) The number of *glaA* mRNA foci in each culture condition. Error bars indicate standard deviation of the mean (n = 3 in D; n = 10 in G). *** Statistically significant difference at P < 0.001. (E) Schematic of transcription inhibition treatment. (F) Colocalization of *glaA* mRNAs (green) and nuclei (magenta) in cells treated with DMSO or ActD. White arrows indicate the hyphal tip. Scale bars, 10 µm.

Next, we investigated whether the dot-like fluorescence visualized by the MS2 system reflected the transcriptional response system of the *glaA* gene. *glaA* transcription is strongly repressed in the presence of glucose and highly activated in the presence of maltose and starch.^47,49^ Therefore, we measured the number of cytoplasmic dot-like fluorescence in glucose and maltose culture conditions using the strain expressing NLS tagged with the red fluorescent protein mCherry to label nuclei. The dot-like fluorescence in the cytoplasm was rarely observed in glucose cultures, while they were frequently observed in the cytoplasm of maltose cultures (Figures 2C and 2D), indicating that the MS2 system guarantees the transcriptional response system of the *glaA* gene.

As further support, the transcriptional response of *glaA* mRNA was analyzed upon the addition of the transcriptional inhibitor actinomycin D (Act D). Using a strain expressing cytoplasmic EGFP under the *amyB* promoter that exhibits a carbon source expression response similar to *glaA*, we confirmed that 100 µg/ml Act D exhibits transcriptional inhibition (Figures S2A-S2C). The *glaA*-MS2 strain was pre-cultured in glucose medium supplemented with Act D for 1 h and then cultured in maltose medium supplemented with Act D for 2 h, and the number of *glaA* mRNA foci in each phase was measured (Figure 2E). The dot-like fluorescence was almost completely absent when Act D was added, suggesting that the dot-like fluorescence in the cytoplasm is derived from transcription (Figures 2F and 2G). Furthermore, when cultured again in maltose medium without Act D, *glaA* transcription resumed and dot-like fluorescence appeared in the cytoplasm (Figures 2F and 2G). Taken together, these results strongly suggest that the dot-like fluorescence visualized by the MS2 system is mRNA-derived fluorescence that depends on the *glaA* mRNA transcriptional response system.

### Localization and dynamics of *glaA* mRNA

Filamentous fungi have an elongated cellular form consisting of a series of hyphal cells with multiple nuclei. Moreover, the cells are numerous and branched, with complex and large cellular spaces. In such multinucleated and multicellular hyphae, we analyzed the localization of mRNA encoding GlaA, a secreted enzyme (Figures 3A and S3A). After culturing cells overnight in the maltose-containing medium to induce *glaA* mRNA expression, hyphal cells were compartmentalized into basal, septum and apical regions, and *glaA* mRNA was found to most prominently localized to the apical region (Figure 3B and S3B). smFISH also confirmed a similar localization pattern of *glaA* mRNA (Figure S3A). In addition, *glaA* mRNA was observed to localize in the vicinity of the septum. Therefore, we compared the number of *glaA* mRNA foci in the middle part of the hyphae between cell compartments with and without septa, and found that *glaA* mRNAs were significantly more abundant in the vicinity of the septa (Figure 3C). The localization of secretory α-amylase at the septa has suggested the possibility of enzyme secretion from the septa,^45^ and the results of *glaA* mRNA localization in the present study may further support this hypothesis. Even at the basal region, where *glaA* mRNA is scarce, it is localized when a septum or a branched hypha is formed (Figures 3D and S3C). This suggests that cellular recognition on the location of the hyphal tip and septum may play an important role in the localization of *glaA* mRNA.

**Figure 3.**
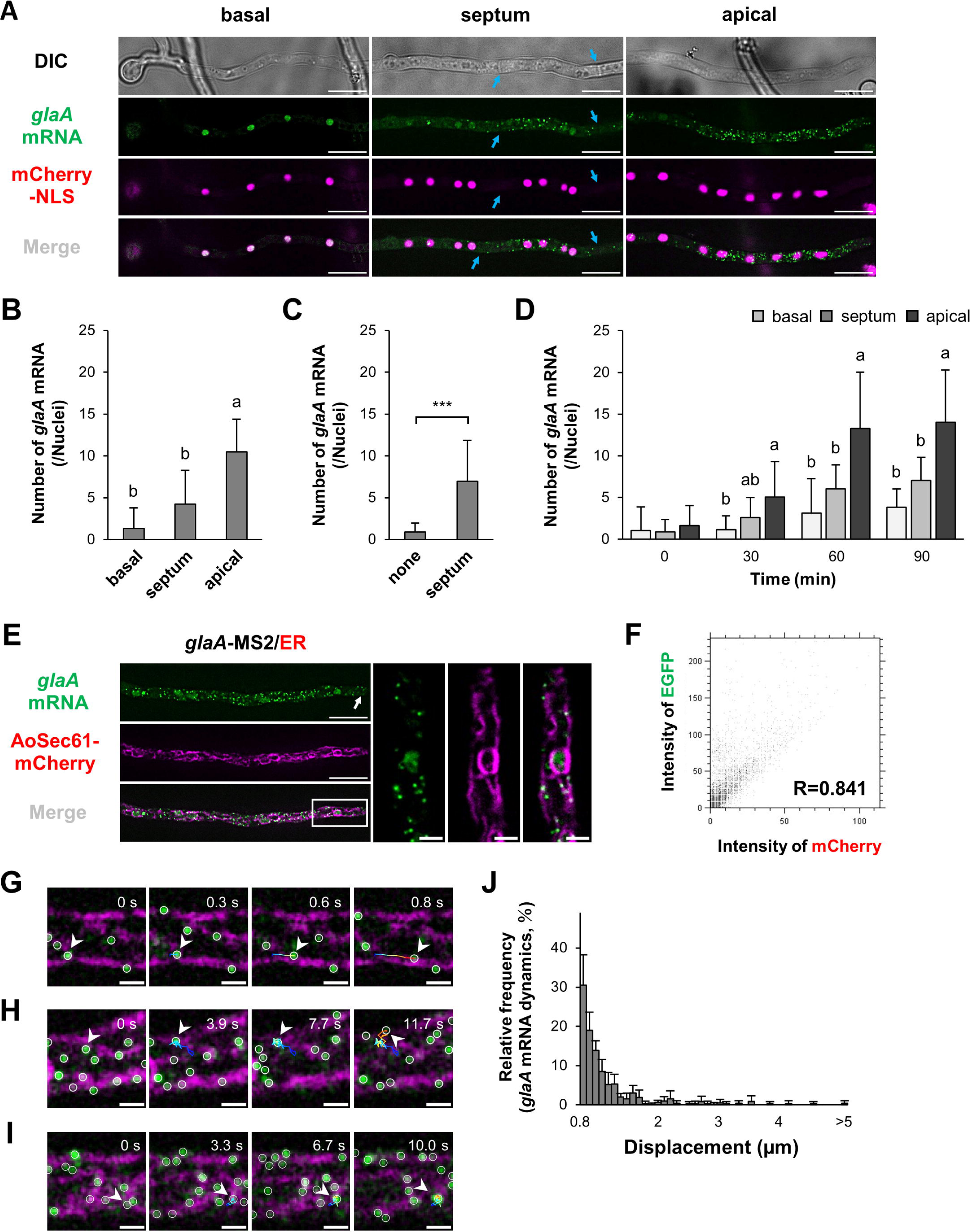
Localization and dynamics of *glaA* mRNA. (A) Colocalization of *glaA* mRNAs (green) and nuclei (magenta). Blue arrows show septa. Scale bars, 10 µm. (B, C and D) The number of *glaA* mRNA foci in each hyphal area and timing after shifting medium. Error bars indicate the standard deviation of the mean (n = 10). Significant difference at P < 0.05 between a and b (Tukey–Kramer test), and *** statistically significant difference at P < 0.001. (E) Colocalization of *glaA* mRNAs (green) and ER (magenta). The white arrow shows the hyphal tip. Scale bars, 10 µm and 2 µm (enlarged images). (F) Colocalization analysis of *glaA* mRNAs (EGFP) and ER (mCherry). (G, H and I) Co-dynamics of *glaA* mRNAs (green) and ER (magenta).White arrowheads show each *glaA* mRNA dynamics. Scale bars, 1 µm. (J) Relative frequency on the displacement of *glaA* mRNA dynamics at the apical region. Error bars indicate the standard deviation of the mean (n = 5).

Next, we analyzed how *glaA* mRNA localizes in the apical and septum regions. Transcription induction of *glaA* mRNA was performed in the maltose-containing medium after incubation in the glucose-containing medium, and the number of *glaA* mRNA foci was measured over time (Figure 3D and S3D). The results showed that *glaA* mRNA was not transcriptionally induced in all nuclei, but was selectively induced in nuclei near the hyphal tip and septum. Therefore, our results suggest that *A. oryzae* can efficiently localize mRNA of a secretory enzyme to specific cellular compartments by selectively activating the transcription of the secretory enzyme gene in the nuclei near the hyphal tip and septum.

Secreted enzymes and plasma membrane proteins have an ER localization signal at their N-termini and are recruited to translational sites on the ER membrane. To analyze the co-localization of *glaA* mRNA and ER, we visualized the ER by fusing mCherry to AoSec61 (AO090701000730) (Figures S3E), a translocon complex component localized at the translocation site in the ER, and found that *glaA* mRNA co-localized with ER with a high degree of correlation (Figures 3E, 3F, and S3F). If *glaA* mRNA is free in the cytoplasm then we would expect there should be a lot of *glaA* mRNA foci independent of the ER, so we analyzed the relationship between *glaA* mRNA dynamics and the ER. *glaA* mRNA exhibited directed dynamics, Brownian-like dynamics and static states, with most dynamics observed on the ER membrane (Figures 3G-3I, Movie S1). Among them, Brownian-like dynamics and static states were the majority, and directional dynamics were present in less than 5% of the cases. (Figure 3J). These results suggest that *glaA* mRNA localization is strongly correlated with the ER and that its dynamics follow the ER structure.

### *glaA* mRNA is transported in the vicinity of ER by an EE-independent manner

The directional dynamics observed in *glaA* mRNA could be attributed to a molecular motor responsible for material transport on the cytoskeleton. Kinesin-1 and kinesin-3, which are responsible for material transport on microtubules, move at an average speed of 0.8∼1 µm/s and 1∼2 µm/s, respectively.^59^ It is unclear whether kinesin-1 is similar to the average rate of kinesin motors in filamentous fungal cells, but analysis of EE dynamics suggests that kinesin-3 has similar kinetic characteristics.^60-63^ Previous studies in various species suggest that kinesin-1 and kinesin-3 on microtubules and myosin-V on actin filaments are responsible for mRNA transport.^6^ Based on these findings, we speculated that the directional dynamics of *glaA* mRNA seen near the ER involves a molecular motor that moves on the cytoskeleton.

In the filamentous fungus *Aspergillus nidulans*, the ER structure of the hyphal tip is suggested to be polarized depending on actin cables rather than microtubules.^64^ In *A. oryzae* we found that ER structures within 50∼70 µm of the hyphal tip are associated with the subcellular localization of *glaA* mRNA (Figure 3E), but the direct relationship between ER structures and mRNAs encoding secretory proteins in that region is not known. Therefore, we analyzed the effects of the cytoskeleton on *glaA* mRNA dynamics using the microtubule polymerization inhibitor nocodazole (Noc) and the actin polymerization inhibitor latrunculin B (Lat B), and evaluated their effects on ER structure. First, we confirmed that microtubule-dependent EE dynamics are no longer observed after Noc treatment and that actin-related protein localization is no longer observed after Lat B treatment (Figures S4A-S4D). However, it was difficult to analyze the dynamics of *glaA* mRNA because its stability was significantly reduced by the addition of Noc and Lat B, with almost complete loss of *glaA* mRNA in the cytoplasm (Figure 4A). On the other hand, the ER structure was not affected by the addition of Lat B, but the reticular structure of the cytoplasmic ER was significantly disrupted by the addition of Noc (Figures 4A-4C), suggesting that the ER structure is formed in a microtubule-dependent manner, similar to what has been reported for microtubule-dependent ER formation in mammalian cells.^65^ These results suggest that the ER structure is dependent on microtubules in *A. oryzae* and that *glaA* mRNA may therefore be transported by molecular motors on microtubules in the vicinity of the ER.

**Figure 4.**
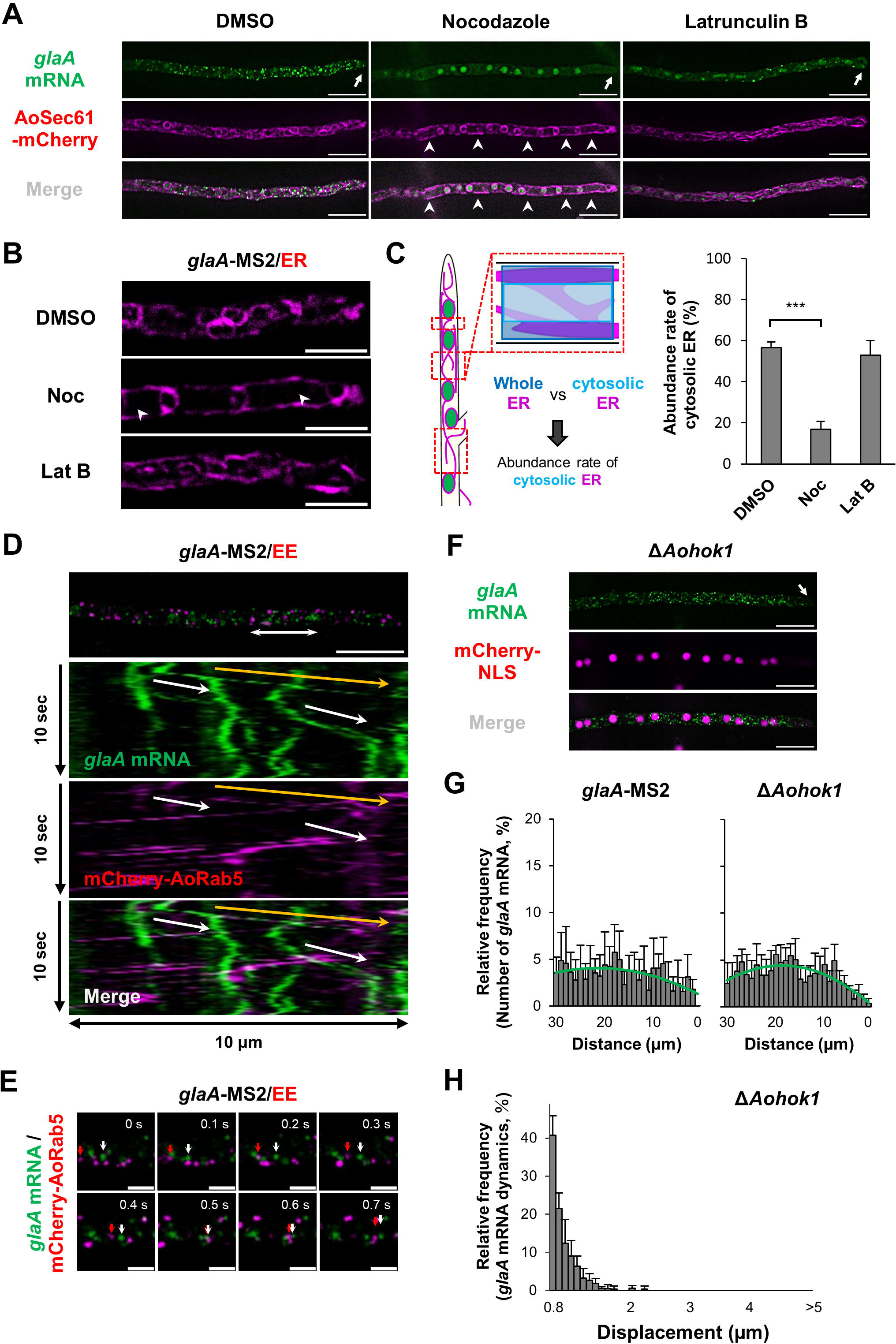
EE-independent *glaA* mRNA motility on microtubule. (A) Colocalization of *glaA* mRNAs (green) and ER (magenta). White arrows show the hyphal tips, and arrowheads show collapsed sites of ER. Scale bars, 10 µm. (B) Enlarged images from (A) at the hyphal tip region in each culture condition. White arrowheads show collapsed sites of ER. Scale bars, 5 µm. (C) Schematic shows how to calculate the abundance rate of cytosolic ER in each culture condition. Error bars indicate the standard deviation of the mean (n = 3). *** Statistically significant difference at P < 0.001. (D) Kymograph of co-dynamics of *glaA* mRNA and EE (mCherry-AoRab5). The white double arrow shows the analyzed region. In kymographs, white arrows show *glaA* mRNA dynamics and the orange arrow shows the time different trajectory of *glaA* mRNA and EE. Scale bar, 10 µm. (E) Time-lapse images of *glaA* mRNA and EE dynamics taken from the orange arrow trajectory in (D). White and red arrows show *glaA* mRNA and EE, respectively. Scale bars, 2 µm. (F) Colocalization of *glaA* mRNAs (green) and nucleus (magenta) in the Δ*Aohok1* strain. The white arrow shows the hyphal tip. Scale bars, 10 µm. (G) Relative frequency of the number of *glaA* mRNA foci at the apical region in *glaA*-MS2 and Δ*Aohok1* strains. Green curves indicate polynomial approximation. Error bars indicate the standard deviation of the mean (n = 5). (H) Relative frequency on the displacement of *glaA* mRNA dynamics at the apical region in the Δ*Aohok1* strain. Error bars indicate the standard deviation of the mean (n = 5).

In *U. maydis*, mRNA is bound to the EE membrane via RNA-binding proteins and other complexes, and is transported through the cell by the microtubule motor proteins kinesin and dynein.^3^ Therefore, we visualized EE by fusing mCherry to AoRab5 and analyzed its co-movement with *glaA* mRNA. In these experiments *glaA* mRNA directional dynamics showed no correlation with the movement of EE (Figure 4D, Movie S2). However, tracking analysis showed that *glaA* mRNA and EE showed dynamics on the same rails at different times (Figure 4E), suggesting that *glaA* mRNA is intracellularly transported in an EE-independent manner. Since *glaA* mRNA does not appear to be co-transported along with EE, we analyzed *glaA* mRNA dynamics in a strain in which EE dynamics were abolished by disrupting AoHok1, a linker between EE and kinesin-3.^46^ The results showed that disruption of *Aohok1* had little effect on *glaA* mRNA localization, but long-range dynamics of *glaA* mRNA were not observed (Figures 4F-4H). In *U. maydis*, *hok1* disruption has been reported to reduce the motility of kinesin-3,^66^ and thus the reduced long-range dynamics of *glaA* mRNA in the *Aohok1* disrupted cells could be due to reduced kinesin-3 functionality.

### Localization mechanism of *glaA* mRNA via kinesin-1 and kinesin-3

After mRNA is transcribed in the nucleus, it is released to the cytoplasm. If *glaA* mRNA is transported in a microtubule molecular motor-dependent manner, it is likely to be transported by kinesin-1 or kinesin-3 to the plus end of microtubules.^63,67-69^ Thus, we analyzed the subcellular localization of *glaA* mRNA upon disruption of kinesin-1 and kinesin-3, AoKin1 (AO090103000408) and AoKin3 (AO090026000806), in *A. oryzae*, respectively (Figures S5A and S5B).

First, we found that EE dynamics almost stopped in Δ*Aokin1* and Δ*Aokin3* cells (Figures S5C and S5D). These results were similar to the EE dynamics when KinA (kinesin-1) and UncA (kinesin-3) were disrupted in *A. nidulans*.^67,68^ Next, we investigated the subcellular localization of *glaA* mRNA, and found that little effect was seen in Δ*Aokin3* cells, whereas almost no localization of *glaA* mRNA was seen at the hyphal tip of Δ*Aokin1* and Δ*Aokin1*Δ*Aokin3* (ΔΔ*k1k3*) cells (Figures 5A-5C). Since *glaA* mRNA localization is regulated by the nuclear location where it is transcribed, it is possible that the effect of *Aokin1* knockout could be due to changes in nuclear localization. Indeed, analysis of the distance from the hyphal tip to the first nucleus revealed that the nucleus was significantly retracted from the hyphal tip in Δ*Aokin1* and ΔΔ*k1k3* cells (Figure 5D). These results suggest that AoKin1 contributes to the proper positioning of nuclei at the apical region and that such a regulatory mechanism is important for the localization of *glaA* mRNA near the secretion sites.

**Figure 5.**
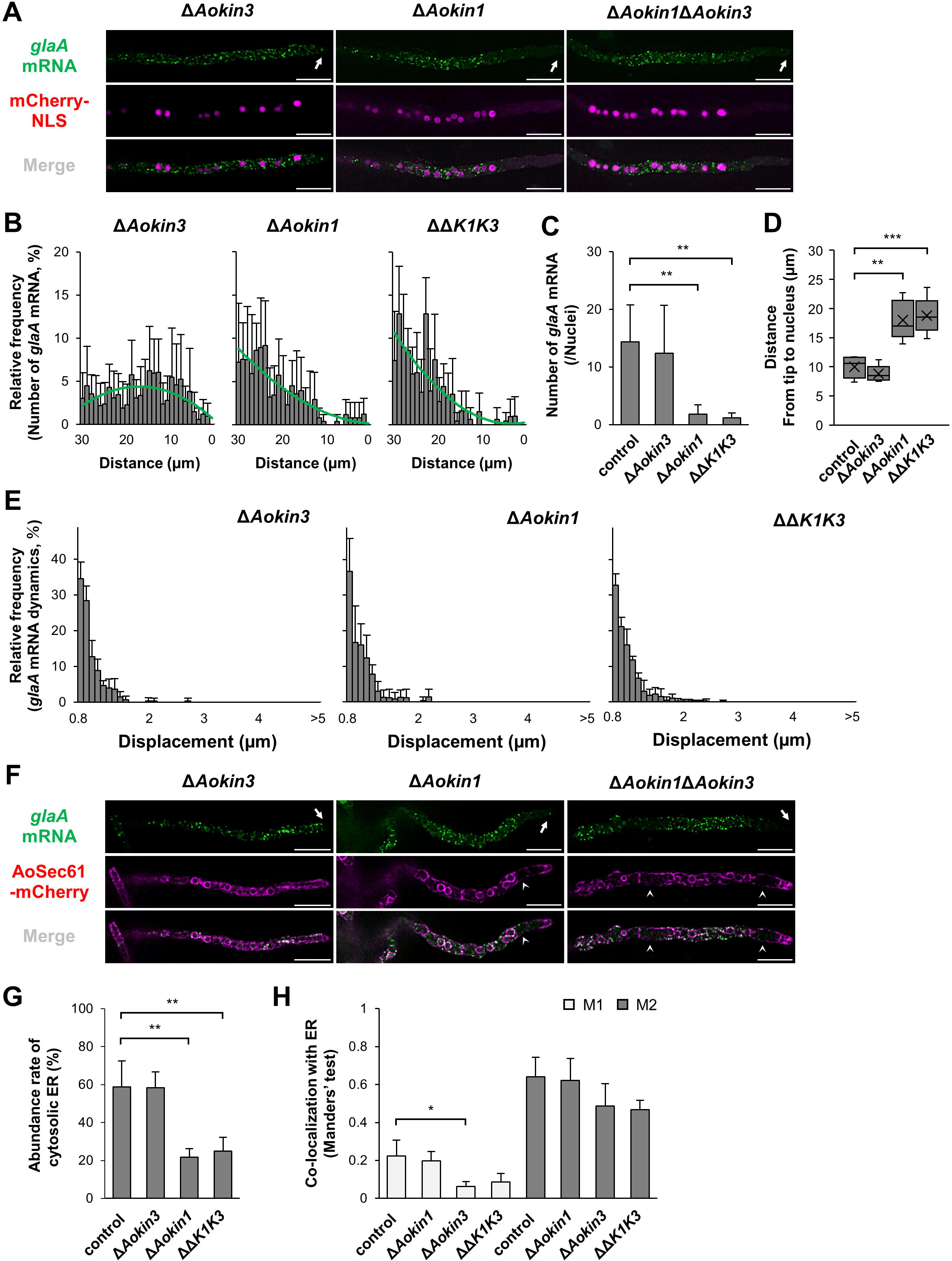
Long-distance *glaA* mRNA motility by kinesin motors. (A) Colocalization of *glaA* mRNAs (green) and nucleus (magenta) in Δ*Aokin1*, Δ*Aokin3* and Δ*Aokin1*Δ*Aokin3* strains. White arrows show the hyphal tips. Scale bars 10 µm. (B) Relative frequency of the number of *glaA* mRNA foci at the apical region in Δ*Aokin1*, Δ*Aokin3* and Δ*Aokin1*Δ*Aokin3* strains. Green curves indicate polynomial approximation. Error bars indicate the standard deviation of the mean (n = 5). (C) The number of *glaA* mRNA foci in the control, Δ*Aokin1*, Δ*Aokin3* and Δ*Aokin1*Δ*Aokin3* strains. Error bars indicate the standard deviation of the mean (n = 5). ** Statistically significant difference at P < 0.01. (D) The distance from the hyphal tip to the first nucleus in the control, Δ*Aokin1*, Δ*Aokin3* and Δ*Aokin1*Δ*Aokin3* strains. Cross marks, top bars and bottom bars indicate the average, the maximum and the minimum distance, respectively (n = 5). ** and *** Statistically significant difference at P < 0.01 and P < 0.001, respectively. (E) Relative frequency on the displacement of *glaA* mRNA dynamics at the apical region in the Δ*Aokin1*, Δ*Aokin3* and Δ*Aokin1*Δ*Aokin3* strains. Error bars indicate the standard deviation of the mean (n = 5). (F) Colocalization of *glaA* mRNAs (green) and ER (magenta) in Δ*Aokin1*, Δ*Aokin3* and Δ*Aokin1*Δ*Aokin3* strains. White arrows show the hyphal tip and white arrowheads show collapsed sites of ER. Scale bars, 10 µm. (G) Abundance rate of cytosolic ER in the control, Δ*Aokin1*, Δ*Aokin3* and Δ*Aokin1*Δ*Aokin3* strains. Error bars indicate the standard deviation of the mean (n = 3). ** Statistically significant difference at P < 0.01. (H) Mander’s test between *glaA* mRNA and ER in the control, Δ*Aokin1*, Δ*Aokin3* and Δ*Aokin1*Δ*Aokin3* strains. M1 indicates ER vs *glaA* mRNA and ER, and M2 indicates *glaA* mRNA vs *glaA* mRNA and ER. Error bars indicate the standard deviation of the mean (n = 3). * Statistically significant difference at P < 0.05.

Next, we analyzed the effect on *glaA* mRNA dynamics in these kinesin disruptants and found that these showed decreased migration distance in all disruptants (Figure 5E). Together with the fact that *glaA* mRNA was transported on the same microtubule rails as EE, this suggests that *glaA* mRNA is transported long distances by AoKin3, although the subcellular distribution of *glaA* mRNA was not significantly affected in Δ*Aokin3* cells (Figure 5B). Moreover, the transport distance of *glaA* mRNA was reduced in Δ*Aokin1* and ΔΔ*k1k3* cells, suggesting that AoKin1 may also be used for long-distance transport.

In animal cells, kinesin-1 contributes to ER elongation and supports the formation of complex ER reticular structures.^70^ In addition, some species have systems that utilize kinesin-1 to transport mRNA.^71^ If kinesin-1 is required for *glaA* mRNA transport, it could be taking place in the vicinity of ER, but since there are no reports of a link between kinesin-1 and ER in filamentous fungi, further analysis of *glaA* mRNA and ER was conducted. The results showed that the ER structure was partially disrupted in Δ*Aokin1* and ΔΔ*k1k3* cells, making it difficult to form a reticular structure (Figures 5F and 5G), suggesting that AoKin1 is partly responsible for the formation of ER reticular structures in filamentous fungi. We analyzed whether such deletion of *Aokin1* affected the localization of *glaA* mRNA to the ER, and found that *glaA* mRNA still co-localized with the ER (Figure 5H). Therefore, it is possible that *glaA* mRNA was recruited from the cytoplasm to the ER by the SPR pathway independent of kinesin-1 and kinesin-3.

At the end of the analysis of kinesin-disrupted strains, the effects on cell growth were examined. Deletion of the amylase gene in *A. oryzae* causes a slight growth retardation on the maltose medium.^72^ However, kinesin disruptants did not exhibit repressed cell growth in all culture conditions tested (Figures S5E-S5I), suggesting that kinesin-mediated *glaA* mRNA dynamics do not affect growth levels. Collectively, these results suggest that *glaA* mRNA is localized to the ER by the SPR pathway even in the absence of kinesin motors, and that sufficient amounts of protein are translated and secreted for growth.

### *glaA* mRNA is regulated by stress granule and processing body

*A. oryzae* is normally cultured at 30°C, but when used in the fermentation and brewing industry, it is sometimes grown at temperatures above 40°C, in which case the cells are considered to be exposed to high temperature stress. In addition, *A. oryzae* has a high protein secretion ability, but large amounts of proteins are translated and present in the ER, causing ER stress.^73^ Thus, it is important to elucidate the regulatory mechanism of secretory enzyme mRNA during such stress and to understand the protein secretion process of *A. oryzae* in detail for further effective industrial utilization. Therefore, we analyzed *glaA* mRNA localization under stress conditions in relation to PB and SG, which are important regulators of mRNA translation and degradation.

We fused mCherry to AoEdc3 (AO090120000481) and AoBre5 (AO090102000619) to visualize PB and SG (Figures S6A and S6B), respectively. Although Bre5 has been identified as a component of SG in budding yeast,^74^ its orthologs in filamentous fungi have rarely been analyzed. AoBre5 is a homolog of G3BP1, which is often used in animal cells for SG analysis, and G3BP1 is a major factor supporting SG formation.^75^ AoPab1, a poly-A binding protein, has been previously analyzed as a SG marker in *A. oryzae*,^76^ and we confirmed that the localization of AoPab1 and AoBre5 showed the same behavior under high temperature stress at 45°C for 10 min (Figures 6A and S6C). Edc3 is a common PB marker widely used from budding yeast to animal cells and plays an important role in PB formation. AoDcp2, a decapping protein, has been previously analyzed as a PB marker in *A. oryzae*, and we confirmed that the localization of AoDcp2 and AoEdc3 shows the same behavior (Figures 6B and S6D). Localization analysis of *glaA* mRNA after incubating cells at 45°C for 10 min showed co-localization with SG and PB (Figures 6A and 6B). Consistent with the previous report,^76^ SG visualized by AoBre5 and AoPab1 was formed at the hyphal tip and co-localized with *glaA* mRNA (Figures 6A and S6C). PBs visualized by AoEdc3 and AoDcp2 also co-localized with *glaA* mRNA under both normal and heat stress condition (Figures 6B and S6D).

**Figure 6.**
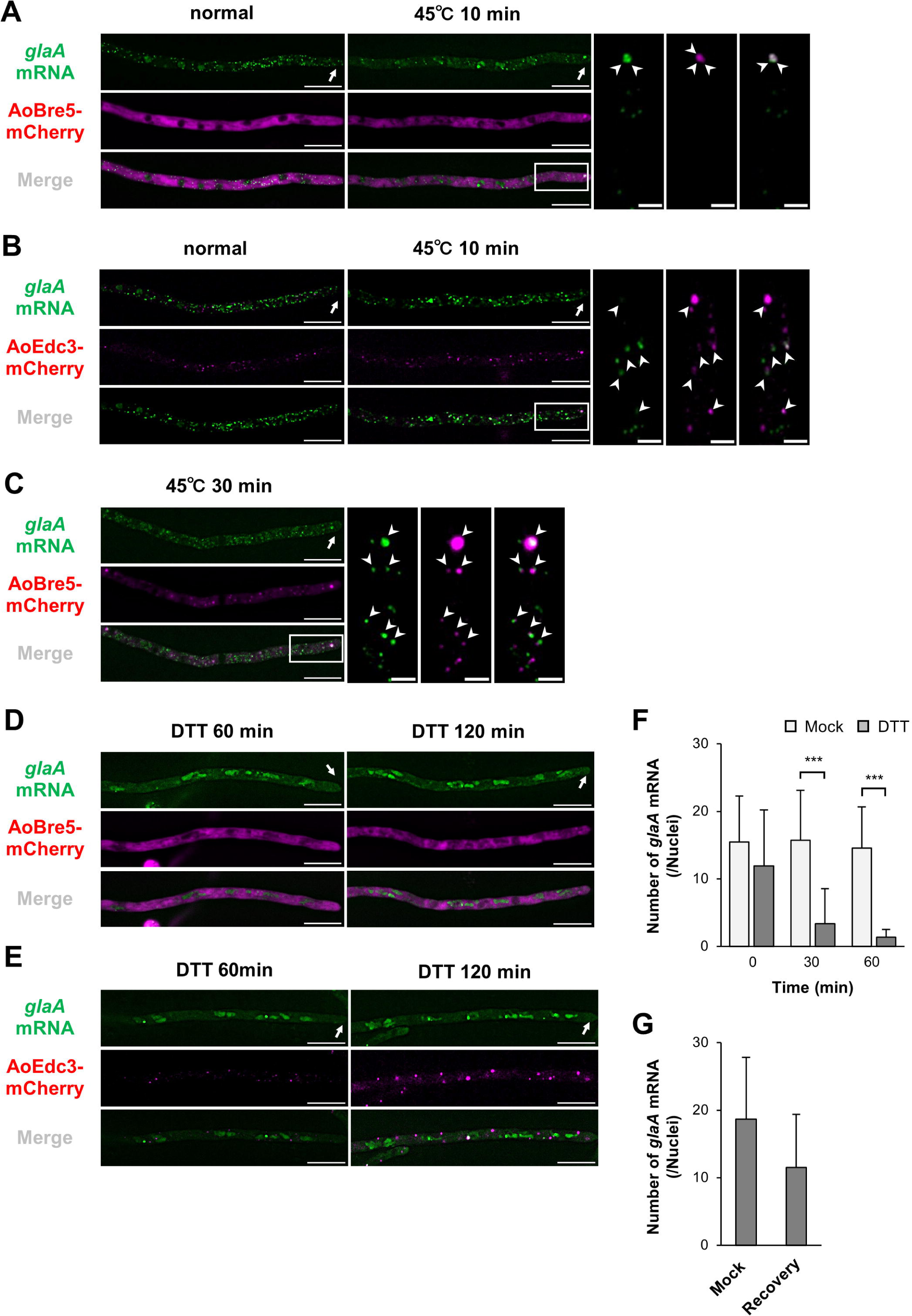
Colocalization of *glaA* mRNA, SG and PB under stress conditions. (A, B and C) Colocalization of *glaA* mRNAs (green) and SG (AoBre5-mCherrry, magenta) or PB (AoEdc3-mCherry, magenta). White arrows show the hyphal tip and arrowheads show colocalization sites. Scale bars, 10 µm and 2 µm (enlarged images). (D and E) Colocalization of *glaA* mRNAs (green) and SG (AoBre5-mCherry, magenta) or PB (AoEdc3-mCherry, magenta). White arrows show the hyphal tip. Scale bars, 10 µm. (F and G) The number of *glaA* mRNA foci in each culture condition. In (G), after 1 h DTT treatment, DTT was washed out and a further 2 h culturing in the fresh medium was performed to count the number of *glaA* mRNAs. Error bars indicate the standard deviation of the mean (n = 10). *** Statistically significant difference at P < 0.001.

Although SGs are important in regulating mRNA localization and translation under stress conditions, they were not expected to form only at the hyphal tips, so the analysis was conducted by exposing the cells to high temperature stress for a longer period of time. In fact, SGs were formed throughout the hyphal cell at 45°C for 30 min, and *glaA* mRNA co-localized with SGs formed at sites other than the hyphal tip (Figure 6C). The co-localization of PBs with SGs formed at 45°C for 30 min (Figure S6E) suggests that SGs formed in the later stage also have components shared with PBs.

When analyzed during ER stress with dithiothreitol (DTT) in addition to high temperature stress,^76^ cytoplasmic *glaA* mRNA rapidly disappeared and no co-localization with PBs or SGs was observed (Figures 6D-6G). SG formation was much slower during DTT treatment, unlike high temperature stress, and only a few hyphae formed SGs after 2 h of DTT treatment (Figure 6D). In contrast, PB increased in size in all hyphal cells after 2 h of DTT treatment, but co-localization with *glaA* mRNA was hardly observed (Figure 6E). Thus, it is suggested that *glaA* mRNA is largely unregulated by SG and PB during ER stress.

## DISCUSSION

Secretory enzymes are transported to outside cells from specific sites depending on the polarity of the cell type. Such a secretory system is a widely conserved process from fungi to animal cells, and is important not only for normal growth but also for adaptation to the ever-changing external environment. In *A. oryzae*, which is one of the safest and best high-secretory protein producers among industrial filamentous fungi, we investigated the subcellular localization and dynamics of *glaA* mRNA, which encodes a secretory glucoamylase. By analyzing *glaA* mRNA in living *A. oryzae* cells using the MS2 system, we found that the *glaA* mRNA is compartmentally localized near secretory sites such as hyphal tips and septa and that *glaA* mRNA exhibits molecular motor-dependent dynamics. Furthermore, we found that *glaA* mRNA is partially colocalized in SG and PB during high temperature stress.

The localization of *glaA* mRNA at the hyphal tip and septum regions suggested that such spatial regulation may maximize the efficiency of transcription, translation and secretory transport (Figures 3A-3C). In particular, the hyphal tip is a highly concentrated area of secretory vesicles, supporting the idea that the hyphal tip is the main site of secretion. More importantly, we found that nuclei located near the secretory sites selectively activate *glaA* transcription in the compartmentalization of its mRNAs in the elongated cell space of filamentous fungi (Figures 3D and S3D). The mechanism of *glaA* mRNA localization by transcriptional regulation may be related to the subcellular localization of a transcriptional activator AmyR and a repressor CreA. The first possible mechanism of transcriptional regulation that can be inferred from the present results is that AmyR may be localized particularly in the nucleus close to the secretory sites, while CreA is not. However, these proteins have previously been reported to localize to all nuclei in the hyphal cells.^51,52^ Nevertheless, it is questionable whether those reports reflected the *bona fide* localization control of AmyR and CreA, since they showed ectopically expressed AmyR and CreA localization. Therefore, real-time localization analysis of endogenous AmyR and CreA is necessary to elucidate the mechanism of *glaA* mRNA transcriptional regulation in detail.

Since most intracellular *glaA* mRNA localized to the ER and some showed dynamics, it was inferred that translational efficiency on the ER membrane is enhanced by transporting *glaA* mRNA near the ER (Figures 3E-3I). The observed *glaA* mRNAs showed directional dynamics, Brownian-like dynamics, and quiescence, most of which were Brownian-like dynamics or quiescence (Figure 3J). It was hypothesized that quiescent *glaA* mRNA is translated on the ER membrane and that other dynamics in the vicinity of the ER may be due to facilitating the diffusion of *glaA* mRNA inside the cell. In *S. cerevisiae*, She2 has been suggested to connect mRNA and ER, but in filamentous fungi there are no orthologs that show high homology to She2.^77^ The subcellular localization of *glaA* mRNA was restricted near the hyphal tip region, especially around 50∼70 µm, but the fact that it was not dispersed throughout the hyphal cell may be related to the ER structure. Indeed, filamentous fungi in general have a highly polarized ER structure in the apical region,^64^ and our results also show that the ER network structure appears to be more complex at 50-70 µm from the hyphal tip than at the base region in *A. oryzae* (Figures 3E and S3D). In *Xenopus laevis*, polarized ER structures have been suggested to contribute to mRNA asymmetry,^78^ and the spatial molecular control mechanisms of ER structure and mRNAs encoding secretory enzymes require further analysis.

We predicted that kinesin-1 would be involved in the dynamics of *glaA* mRNA near the ER and that kinesin-3 would be involved in its long-distance dynamics (Figure 7A). Even with the deletion of Myo4, the molecular motor that transports *ASH1* mRNA in *S. cerevisiae*, *ASH1* mRNA is still transported on the ER structure.^77^ Similarly, long-distance transport of *glaA* mRNA is no longer observed in the *Aokin1* disruptant, but localization to the ER membrane is still observed. However, in order to determine at what specific stage kinesin-1 is involved in the transport of *glaA* mRNA, it is necessary to visualize kinesin-1 as a single molecule and analyze its co-localization with *glaA* mRNA. In addition, it was an unexpected finding that kinesin-3 was involved in the long-range dynamics of *glaA* mRNA, but in an EE-independent manner (Figures 4D and 4E). Previous models of mRNA transport in filamentous fungi, particularly in *U. maydis*, have suggested that it is EE-dependent,^3^ suggesting the existence of a different mechanism than previously thought for mRNA transport or potentially parallel mechanisms depending on specific mRNAs. For example, the mechanism of mRNA transport that does not utilize membrane transport in other species suggests that mRNA may be transported by kinesin-3 via unidentified RNP granules or complexes.^6^ However, the *Aokin3* disruptant did not significantly affect the subcellular localization of *glaA* mRNA, suggesting that long-distance transport of secretory enzyme mRNA by kinesin-3 is only to support diffusion at the ER membrane and is not an essential mechanism for the subcellular localization of *glaA* mRNA. On the other hand, analysis of the *Aokin1* disruptant, in which nuclei are retracted from the hyphal tip, suggests that proper nuclear positioning is important for the subcellular localization of *glaA* mRNA.

**Figure 7.**
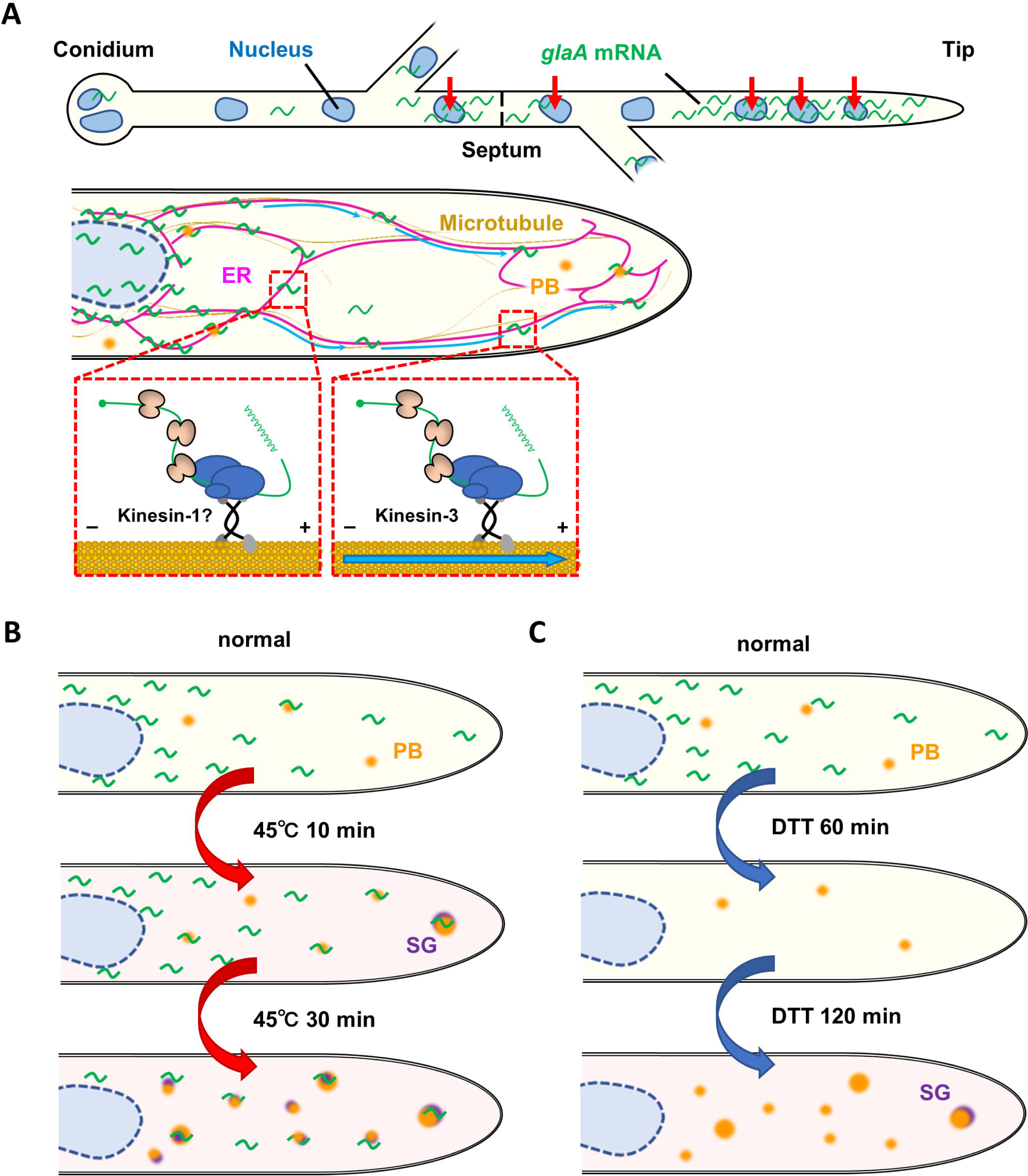
Schematic of localization mechanisms of *glaA* mRNA. (A) The top of the schematic indicates localization of *glaA* mRNA by spatiotemporal transcriptional regulation. Red arrows indicate transcriptional nuclei. The bottom of the schematic indicates *glaA* mRNA dynamics by kinesin motors on the microtubule in the vicinity of the ER membrane. Blue arrows indicate long-directed *glaA* mRNA dynamics. (B and C) Schematics of localization mechanisms of *glaA* mRNAs, PB and SG under heat (B) and ER stresses (C).

Under high temperature stress conditions, *glaA* mRNA localized to SG and PB, suggesting that translation regulation of secretory enzyme mRNA may be one means of rapid environmental adaptation during stress (Figures 6A and 6B). Exposure to longer high temperature stress caused SGs to form throughout the hyphal cell and co-localize with *glaA* mRNA (Figure 6C). The hyphal tip, where the cell wall is thinner and less rigid than in other parts of the hyphal cell, may be able to sense environmental stress more rapidly.^79^ It was also suggested that the formation of SGs is divided into two phases, with the type of stress and exposure time controlling the rate and amount of their formation (Figure 7B). In *A. oryzae*, SG formation is first induced at the hyphal tip under other stresses than high temperature, so the pattern of SG formation may be common under various conditions.^76^ In general, the size and number of SGs vary with the type and duration of stress, and they aggregate significantly when exposed to longer periods of stress.^80^

Although this study showed that secretory enzyme mRNAs co-localize with SGs, mRNAs in SGs generally contain few ER-localized mRNAs.^81^ On the other hand, some reports indicate that ER-localized mRNAs are incorporated into SGs when DTT is present, leading some to believe that the response depends on the type of ER-localized mRNA, and the relationship between ER-localized mRNA and SGs remains unclear.^37,82^ In this study, *glaA* mRNA was incorporated into SGs during high temperature stress, but was rapidly degraded and not incorporated into SGs during ER stress caused by DTT (Figures 6D-6G). Thus, it is suggested that the regulation of ER-localized mRNAs may differ between environmental stresses such as temperature and intracellularly induced stresses. In Hela cells, SGs were clearly formed in the cells 1 h after the addition of DTT, while in *A. oryzae*, almost no SGs were formed, suggesting that the response to SG formation may differ between animal and fungal cells.^37^ Filamentous fungi are known to experience stress when they hyper-secrete proteins, which causes feedback repression of transcription of genes encoding secretory proteins, known as repression under secretion stress (RESS).^83,84^ Thus, it is possible that filamentous fungi, unlike animal cells, have a much more rapid transcriptional repression by RESS during ER stress, and may also have a very high capacity for mRNA degradation. In contrast, unlike SG, PB increased in size during ER stress, but co-localization with *glaA* mRNA was not observed (Figure 7C). Thus, PB may play a more important role as a mRNA regulator of ER stress in filamentous fungal cells.

In this study, we show that in filamentous fungi, mRNA encoding a secretory enzyme is localized to cellular compartments near the site of secretion by spatiotemporal transcriptional regulation, which would support maximizing the flow of transcription, translation, and secretory transport. Moreover, we have elucidated when and where secreted enzyme mRNAs are localized in multinucleated multicellular cells, but it remains an open question as to what transport machinery is responsible for their diffusion on the ER and where they are translated. SunTag and CRISPR-Sunspot have been developed as a means of visualizing the translation process, and SunTag in particular has been widely used from budding yeast to animal cells.^58,85,86^ The application of such techniques to *A. oryzae* will help to elucidate the molecular mechanisms involved in the subcellular localization and translation of secretory enzyme mRNAs. In addition, analysis of the translational dynamics of secretory enzyme mRNAs in relation to SGs and PBs is expected to shed light on the translational regulation of secretory enzyme mRNAs by SGs and PBs. Further understanding of the molecular mechanism will lead to visualization of the rate-limiting step in protein secretion, which will contribute to the molecular breeding of strains capable of higher secretory protein production in the future.

## Supporting information

Resource table

## ACKNOWLEDGEMENTS

We would like to thank Katsuya Gomi for providing the pANPXP-Cre vector, Yoshinori Katakura for supporting the qRT-PCR experiment and Yukie Shima for technical support. We are grateful to the Center for Advanced Instrumental and Educated Supports at the Faculty of Agriculture, Kyushu University for technical help with fluorescence microscopy. This work was supported by JST SPRING grant number JPMJSP2136 to Y.M., JSPS KAKENHI grant number JP22H02245 and NISR Investigator Research Grant to Y.H.

## AUTHOR CONTRIBUTIONS

Y.M. and Y.H. designed and performed experiments. Y.M., K.T., B.M.C. and Y.H. analyzed the data. Y.M., K.T., B.M.C. and Y.H. wrote the manuscript. Y.H. performed all project coordination and secured funding.

## DECLARATION OF INTEREST

The authors declare no competing interests.

## STARfllMETHODS

### Culture media

For selecting *A. oryzae* transformants by using the strain NSPlD1 as a host and the *pyrG* selective marker, M medium (0.2% NH_4_Cl, 0.1% (NH_4_)_2_SO_4_, 0.05% KCl, 0.05% NaCl, 0.1% KH_2_PO_4_, 0.05% MgSO_4_·7H_2_O, 0.002% FeSO_4_·7H_2_O and 2% glucose, pH 5.5) supplemented with 0.15% methionine (MM) was used. In using *niaD* or *sC* selective markers, Czapek-Dox (CD) medium (0.3% NaNO_3_, 0.2% KCl, 0.1% KH_2_PO_4_, 0.05% MgSO_4_·7H_2_O, 0.002% FeSO_4_·7H2O, and 2% glucose, pH 5.5) or CD+0.0015% methionine medium was used according to each strain auxotrophy. When necessary, maltose and potato starch were used as carbon sources. In application of the MS2 system, after inserting a 36× MBS sequence downstream of the *glaA* gene by the Cre/lox system, the cells were cultured on MM medium with the carbon source replaced by 2% xylose to drop off the *pyrG* selection marker and Cre expression cassette. DPY medium (0.5% yeast extract, 1% hipolypeptone, 2% dextrin, 0.5% KH_2_PO_4_, 0.05% MgSO_4_·7H_2_O, pH 5.5) was used as a complete medium.

### DNA cloning

PrimeSTAR Max DNA Polymerase (Takara) and PrimeSTAR GXL DNA Polymerase (Takara) were used to amplify DNA fragments, and In-Fusion HD Cloning Kit (Takara) was used to ligate DNA fragments. In the plasmid generation used in the MS2 system, 24×MBSV6 was amplified from pET264 (Addgene), and the MCP sequence was synthesized with codon-optimization for *A. oryzae* (Thermo Fisher Scientific). The wild-type strain RIB40 was used as a template for the amplification of the *A. oryzae* genomic DNA sequence. We used pANPXP-Cre to insert a 36× MBS sequence downstream of the *glaA* gene by Cre/lox system,^87^ and 939 bp of the 3’ end of the *glaA* ORF and 950 bp downstream were amplified by PCR. First, a NotI restriction enzyme site was inserted upstream of *lox-AnptrA-Cre-lox* by cloning a DNA fragment from inverse PCR of pANPXP-Cre with notI-Fw-1 (GGTTAACGCGGCCGCAGGATTACGTTGAAGGGCTTAG) and notI-Rv-1 (TAATCCTGCGGCCGCGTTAACCGCTCACAATTCCACAC). Then, 36× MBS was amplified with mbs-Fw-1 (GGTTAACGCGGCCGCCAGAGCCCCCTGGCAATCGC), mbs-Rv-1 (GCTATACGAACGGTATCCGTGTGAGGGTCTCTGTCGC), which was cloned into the resultant plasmid that was amplified by inverse PCR with lox-Fw-1 (TACCGTTCGTATAGCATACATTATACGAAG) and amp-Rv-1 (GCGGCCGCGTTAACCGCTCA). Thereafter, the plasmid was amplified by inverse PCR with notI-Fw-2 (AACGGTAGCGGCCGCTATGGTGCACTCTCAGTACAATC) and notI-Rv-2 (CACCATAGCGGCCGCTACCGTTCGTATAATGTATGCTATACG) to introduce a NotI site, resulting in pNotI-36×*mbs*-*lox*-*AnptrA*-*Cre*-*lox*-NotI as a 36×MBS cassette.

The *glaA*-MBS cassette was cloned in three steps. First, the 3’ terminal region 939 bp of *glaA* ORF was amplified with *glaA*-3’U-Fw-1 (GGTTAACGCGGCCGCCGGTCGTGCTGAGAATCAAGCTG) and *glaA*-3’U-Rv-1 (TGCCAGGGGGCTCTGTCACCGCCAAACATCGCTCTG), and cloned into a DNA fragment of 36× MBS cassette that was amplified by inverse PCR with mbs-Fw-2 (CAGAGCCCCCTGGCAATCGC) and amp-Rv-1. Then, 950 bp downstream of *glaA* ORF was amplified with *glaA*-down-Fw-1 (ATTATACGAACGGTATCATCATGTCCCGATGAAGAGG) and *glaA*-down-Rv-1 (CACCATAGCGGCCGCGGAACAACGAGCTTCGGATG), and cloned into the resultant plasmid that was amplified by inverse PCR with amp-Fw-1 (GCGGCCGCTATGGTGCACTC) and lox-Rv-1 (TACCGTTCGTATAATGTATGCTATACGAAGTTATC), resulting in *glaA*-MBS-AnptrA cassette. Finally, the *pyrG* selection marker was amplified with pyrG-Fw-1 (GTATCCAAGACGCATGAGTGCGGCTGACAACTATG) and pyrG-Rv-1 (TCTGACGCCTCATTCCCACGATTGAACATCCTCGTCC), and cloned into the *glaA*-MBS-AnptrA cassette that was amplified by inverse PCR with PxylP-Fw-1 (GAATGAGGCGTCAGACGTGAC) and lox-Rv-2 (ATGCGTCTTGGATACGCGAC) to produce the *glaA*-MBS cassette.

The *pgkA* promoter (P*pgkA*) was used for the expression of MCP in the MS2 system. First, MCP was amplified with mcp-Fw-1 (TTGTGTGATAGAACAATGCTCGCCGTCAAGATGGC) and mcp-Rv-1 (GCCCTTGCTCACCATGTTGAGGCAACGAGAATCGGCGTAGATGCCGCTGTTGGCGG), *egfp* was amplified with egfp-Fw-1 (ATGGTGAGCAAGGGCGAGGA) and egfp-Rv-1 (GAATTATCAACTATGTTACTTGTACAGCTCGTCCATGC), T*amyB*-*niaD*-amp-ori-P*pgkA* was amplified from pgPpgkA-lifeact-egfp-niaD as a template with TamyB-Fw-1 (CATAGTTGATAATTCACTGGCCGTCG) and PpgkA-Rv-1 (TGTTCTATCACACAAGGTGGGGG) and these amplicons were cloned to produce pgPpMG.^88^ Next, *egfp* was amplified with egfp-Fw-2 (TACGCCGATTCTCGTATGGTGAGCAAGGGCGAGGA) and egfp-Rv-2 (CACCATGTTGAGGCACTTGTACAGCTCGTCCATGCCG), pgPpMG was amplified by inverse PCR with egfp-Fw-3 (TGCCTCAACATGGTGAGCAAG) and mcp-Rv-2 (ACGAGAATCGGCGTAGATGC) and these amplicons were ligated to produce pgPpM2G. Then, to incorporate the SV40-derived NLS, P*Aocyc1* (AO090026000267) was amplified from *A. oryzae* RIB40 genomic DNA with PAocyc1-Fw-1 (GATAACAATTTCACACTACTAGGGGGAATACAGACGG) and PAocyc1-Rv-1 (TTACGCTTCTTTTTAGGCATTGTGAATGTGTGTAGTAAAGTCTAAAAGAC), pgPpM2G was amplified by inverse PCR with S40nls-Fw-1 (TAAAAAGAAGCGTAAAGTGACCGGCATGCTCGCCGTCAAGATGGC) and amp-Rv-2 (TGTGAAATTGTTATCCGCTGGTATCAG) and these amplicons were ligated to produce pgPcNM2G. Lastly, P*pgkA* was amplified with PpgkA-Fw-1 (GATAACAATTTCACATATTGACTACTATGGTAACCAACGCG) and PpgkA-Rv-2 (CTTCTTTTTAGGCATTGTTCTATCACACAAGGTGGGG), pgPcNM2G was amplified by inverse PCR with S40nls-Fw-2 (ATGCCTAAAAAGAAGCGTAAAGTGAC) and amp-Rv-2 and these amplicons were ligated to produce pgPpNM2G.

In the preparation of plasmids for co-localization analysis, *Aosec61*, *Aorab5*, *Aobre5*, *Aopab1*, *Aoedc3*, *Aodcp2* and AoStuA-derived NLS were amplified by PCR from *A. oryzae* genomic DNA. *Aosec61* was amplified with S61-Fw-primer (TTGTGTGATAGAACAATGAGCGGACGTGAGTGCAC) and S61-Rw-primer (GCCCTTGCTGACCATGTTGCCAGGAACAAGGCCCTTG), *mcherry* was amplified with mCh-Fw-1 (ATGGTCAGCAAGGGCGAAGAG) and mCh-Rv-1 (GAATTATCAACTATGTTACTTGTACAGCTCATCCATGCC), T*amyB*-*niaD*-amp-ori-P*pgkA* was amplified with TamyB-Fw-1 and PpgkA-Rv-1 and these amplicons were ligated to produce pgPpS61mCN. Then, to replace the *niaD* selection marker, the *sC* selection marker was amplified with sC-Fw-1 (ATTTCACACCGCATAGATTTAGTTCCGTTCGTGCAGG), sC-Rv-1 (TGAGAGTGCACCATACCAAGGAACAGGTCAGGATTAAAG), pgPpS61mCN was amplified by inverse PCR with amp-Fw-2 (TATGGTGCACTCTCAGTACAATCTGC) and TamyB-Rv-1 (TATGCGGTGTGAAATACCGCACAG) and these amplicons were ligated to produce pgPpS61mCS.

*Aorab5* was amplified with R5-Fw-1 (GATGAGCTGTACAAGATGTCTGAGTCAACTAGCACAAATAC) and R5-Rw-1 (GAATTATCAACTATGTTAACAAGCACAACCTTCTTTGGC), *mcherry* was amplified with mCh-Fw-2 (TTGTGTGATAGAACAATGGTCAGCAAGGGCGAAGA) and mCh-Rv-2 (CTTGTACAGCTCATCCATGCCG), T*amyB*-*sC*-amp-ori-P*pgkA* was amplified from pgPpS61mCS with TamyB-Fw-1 and PpgkA-Rv-1 and these amplicons were ligated to produce pgPpmCR5.

*Aobre5* was amplified with B5-Fw-1 (TTGTGTGATAGAACAATGGCCGATACTCAAGCCCCC) and B5-Rw-1 (GCCCTTGCTGACCATAGCAGCCTGAGCCTGGTTGC), *mcherry*-T*amyB*-*sC*-amp-ori-P*pgkA* from pgPpS61mCS was amplified with mCh-Fw-1 and PpgkA-Rv-1 and these amplicons were ligated to produce pgPpB5mC.

*Aoedc3* was amplified with E3-Fw-1 (TTGTGTGATAGAACAATGGATCGCGCCCGCAAGAAG) and E3-Rw-1 (GCCCTTGCTGACCATAGCAGAGGATGGCTGGTAGCG), *mcherry*-T*amyB*-*sC*-amp-ori-P*pgkA* was amplified from pgPpS61mCS with mCh-Fw-1 and PpgkA-Rv-1 and these amplicons were ligated to produce pgPpE3mC.

AoPab1-EGFP and AoPab1-mCherry expression plasmids were prepared as follows. First, P*pgkA* was amplified by PpgkA-Fw-1 and PpgkA-Rv-3 (GCTCACCATCCCGGGTGTTCTATCACACAAGGTGGGGGG) and pgP*amyB*-SmaI-*egfp*-T*amyB* (Morita et al., 2021) was amplified by inverse PCR with egfp-Fw-4 (CCCGGGATGGTGAGCAAGGG) and amp-Rv-2 and these amplicons were ligated to produce pgPpSmaIG. *Aopab1* was amplified with P1-Fw-1 (TGTGATAGAACACCCATGTCTGCCGACGCCTCTAC) and P1-Rv-1 (CTTGCTCACCATCCCCGACTTGTTCTCTTCGGTAGAGG) and cloned into a SmaI digested DNA fragment of pgPpSmaIG, resulting in pgPpP1G. To introduce a SmaI restriction enzyme site upstream of *mcherry*, pgPpS61mCS was amplified by inverse PCR with mCh-Fw-3 (TTGTGTGATAGAACACCCGGGATGGTCAGCAAGGGCGAAGA) and PpgkA-Rv-1 to produce pgPpSmaImC. *Aopab1* was amplified with P1-Fw-1 and P1-Rv-2 (CTTGCTGACCATCCCCGACTTGTTCTCTTCGGTAGAGG) and cloned into a DNA fragment of pgPpSmaImC digested with SmaI to produce pgPpP1mC.

*Aodcp2* was amplified with D2-Fw-1 (TGTGATAGAACACCCATGACAGAAACAAAGATGCATTTAGAAG) and D2-Rw-1 (CTTGCTGACCATCCCCTTGTTCCCTTTGGCAACACC) and cloned into a DNA fragment digested with SmaI of pgPpSmaImC to produce pgPpD2mC.

To generate pgPtmCN, the 1,009 bp promoter region of *Aotps1* (AO090003000417) was amplified from *A. oryzae* RIB40 genomic DNA with Ptps1-Fw-1 (GATAACAATTTCACACGCCACCATGAGAGAAAAGG) and Ptps1-Rv-1 (GCCCTTGCTGACCATTTTTGGTGAAATCGGATTTGATGGTTG), *mcherry* was amplified with mCh-Fw-1 and mCh-Rv-3 (ACGCTTTCCTGACCCCTTGTACAGCTCATCCATGCCG), AoStuA-derived NLS was amplified with SAnls-Fw-1 (GGGTCAGGAAAGCGTATGCGGG) and SAnls-Rw-1 (TTATCAACTATGTTAAGTCTTCCGGCGCTTGCTCT), T*amyB*-*sC*-amp-ori was amplified with TamyB-Fw-2 (TAACATAGTTGATAATTCACTGGCCGTC) and amp-Rv-2 and these amplicons were ligated.

In the preparation of the ΔAohok1-pyrG cassette, 962 bp upstream of the *Aohok1* ORF was amplified with Aohok1-up-Fw-1 (GGTTAACGCGGCCGCCATTCCTAGGGTGGACGGATC) and Aohok1-up-Rv-1 (TTGTCAGCCGCACTCGTTGGCCTAAGAGGCAGTTG), 982 bp downstream of the *Aohok1* ORF was amplified with Aohok1-down-Fw-1 (GATGTTCAATCGTGGTACCGAAGCATAATTTGCAGTATATCC) and Aohok1-down-Rv-1 (CACCATAGCGGCCGCGCCTATATGGATCCGGTGTCG), the *pyrG* selection marker was amplified with pyrG-Fw-2 (GAGTGCGGCTGACAACTATG) and pyrG-Rv-2 (CCACGATTGAACATCCTCGTCC), vector sequences containing amp were amplified with amp-Fw-1 and amp-Rv-1 and these amplicons were ligated.

For the preparation of the ΔAokin1-pyrG cassette, 936 bp upstream of the *Aokin1* ORF was amplified with Aokin1-up-Fw-1 (GGTTAACGCGGCCGCCAAAAGGTGACAGCATACTGCC) and Aokin1-up-Rv-1 (TTGTCAGCCGCACTCGTTCACTCTGACCCAGAAACCC), 956 bp downstream of the *Aokin1* ORF was amplified with Aokin1-down-Fw-1 (GATGTTCAATCGTGGTCCACGATTTCTCTTTTGTTGTTTATTTTATC) and Aokin1-down-Rv-1 (CACCATAGCGGCCGCGAGGCCTTACTGAAGGTGCAG), the *pyrG* selection marker was amplified with pyrG-Fw-2 and pyrG-Rv-2, and vector sequences containing amp were amplified with amp-Fw-1 and amp-Rv-1 and these amplicons were ligated. Then, the *AnptrA* selection marker was amplified with AnptrA-Fw-1 (TGGTCGCGTATCCAAGACGC) and AnptrA-Rv-1 (TCATTCACTTTGTTGCCTCGGTG), the ΔAokin1-pyrG cassette was amplified by inverse PCR with Aokin1-down-Fw-2 (CAACAAAGTGAATGATCCACGATTTCTCTTTTGTTGTTTATTTTATC) and Aokin1-up-Rv-2 (TTGGATACGCGACCAGTTCACTCTGACCCAGAAACCC) and these amplicons were ligated to produce the ΔAokin1-ptrA cassette.

In the preparation of the ΔAokin3-pyrG cassette, 920 bp upstream of the *Aokin3* ORF was amplified with Aokin3-up-Fw-1 (GGTTAACGCGGCCGCGCAAGAGCATTTCACATGGGC) and Aokin3-up-Rv-1 (TTGTCAGCCGCACTCTATGACCAATCCCCCATGCAC), 968 bp downstream of the *Aokin3* ORF was amplified with Aokin3-down-Fw-1 (GATGTTCAATCGTGGCTTCTGTCGTCACATCAGCC) and Aokin3-down-Rv-1 (CACCATAGCGGCCGCCTTCCGGCCATGACCAGTAC), the *pyrG* selection marker was amplified with pyrG-Fw-2 and pyrG-Rv-2, the vector sequence containing amp was amplified with amp-Fw-1 and amp-Rv-1 and these amplicons were ligated.

In the generation of AoAbp1-EGFP expression strain, approximately 1 kb of the ORF of *Aoabp1*, excluding the stop codon, was amplified by PCR with Aoabp1-nsc-att-Spe-F (GGGGACAACTTTGTATAGAAAAGTTGACTAGTAGTAAACGATGCCACGGGCAACAG) and Aoabp1-nsc-att-R (GGGGACTGCTTTTTGTTACACTTGCTCTTCGAAGTTCTACATATTTGCTGGG), about 1 kb downstream of *Aoabp1* was amplified by PCR with Aoabp1-dw-att-F (GGGGACAGCTTTCTTGTACAAAGTGGTCCAAGGTGGTGTCTTCCAC) and Aoabp1-dw-att-Spe-R (GGGGACAACTTTGTATAATAAAGTTGACTAGTGAAGCCACGGCTATTGATCTTG) and these PCR amplicons were cloned as pg5’abp1 and pg3’abp1, respectively. These plasmids were ligated with pgegTasC and pDEST^TM^R4-R3 by the LR reaction of the MultiSite Gateway System.^46,89^ DNA was fragmented by SpeI treatment and transformed into the strain NSRku70-1-1A using selection by the *AosC* marker.^90^

For the complementation of *pyrG*, pgSmaI-pyrG-ptrA was created. The *pyrG* selection marker was amplified from *A. oryzae* genomic DNA and the *ptrA* selection marker was amplified from pPTRII (Takara) by PCR. The *pyrG* selection marker was amplified by PCR with pyrG-Fw-3 (GATAACAATTTCACACCCGGGGAGTGCGGCTGACAACTATG) and pyrG-Rv-3 (CGTAATCAATTGCCCCCACGATTGAACATCCTCGTCC), the *ptrA* selection marker was amplified by PCR with ptrA-Fw-2-primer (GGGCAATTGATTACGGGATCCC) and ptrA-Rv-2 (TGAGAGTGCACCATAATGGGGTGACGATGAGCCGCTCTTG), amp sequence was amplified by PCR with amp-Fw-2 and amp-Rv-2 and these PCR amplicons were ligated, resulting in pgSmaI-pyrG-ptrA. To complement *AosC*, pgegTasC was used to remove sequences of *egfp* and T*amyB*,^46^ resulting in pgNotI-AosC-NotI.

### Growth assay

Growth assay was performed using CD (mal) medium with the carbon source replaced by maltose, CD (PS) medium with the carbon source replaced by potato starch and DPY medium as a complete medium, inoculated with 1.0 × 10= conidia/10 µl and incubated at 30°C for 3 days. Growth assay of each kinesin motor deletion strain was performed using a medium based on CD+0.0015% methionine medium, and colony diameters were measured.

### Quantitative reverse transcription PCR analysis

Quantitative reverse transcription PCR (qRT-PCR) analysis was conducted as described previously.^46^ The NSlDSN1 and *glaA*-MS2 strains were inoculated with 1.0 × 10= of their conidia in 100 ml of CD (mal) medium and cultured at 30°C, 24 h, 150 rpm. Total RNA was then extracted from each mycelium according to the instructions of the RNeasy^®^ Plant Mini Kit (Qiagen), and cDNA was synthesized using the SuperPrep Cell Lysis & RT Kit for qPCR (Toyobo). Thermal Cycler Dice Real Time System TP-800 instrument (Takara) and Thunderbird SYBR qPCR Mix (Toyobo) were used for qRT-PCR analysis of each cDNA. Expression levels of *glaA* mRNA were measured using glaA-Fw-RT (CGGTTCCTCGTGCCCCTATT), glaA-Rv-RT (GATGTCTTTGCCCGATCGCC). Expression of *gpdA* as normalization was measured using gpdA -Fw-RT (CGTCGAGTCCACTGGTGTCTT), gpdA -Rv-RT (TTGTTGACACCCATAACGAACATGG).

### Glucoamylase activity analysis

Glucoamylase activity analysis was performed as described previously.^46^ Twenty ml of DPY medium was inoculated with 1.0 × 10= of the NSlDSN1 and *glaA*-MS2 strains and cultured at 150 rpm at 30°C for 3 days. Each culture supernatant was then collected and glucoamylase activity was measured according to the instructions (Kikkoman).

### Fluorescence microscopy

For smFISH, an ECLIPSE Ti2-A inverted microscope (Nikon) was employed, equipped with a CFI Plan Apo Lambda 100 × objective lens (1.45 numerical aperture), a DS-Qi2 digital camera, an LED-DA/FI/TX-A triple band filter (Semrock: Exciter, FF01-378/474/575; Emission, FF01-432/523/702; Dichroic mirror, FF409/493/596-Di02), an LED light source X-LED1 and differential interference contrast (DIC) to observe the EGFP, CAL Fluor Red 610, and DAPI fluorescent signals and hyphal morphology of *A. oryzae* cells.

Live-cell imaging of *glaA* mRNA by MS2 system was performed using THUNDER Imager Live Cell (Leica microsystems), equipped with an HCX PL APO 100×/1.40-0.70 OIL objective lens, a K8 Scientific CMOS camera, a DFT51010 filter (Leica : LED, DAPI_FITC_TXRED_Cy5-390/475/555/635; Excitation Range, DAPI_FITC_TXRED_Cy5-375-407/462-496/542-566/622-654; Emission Range, DAPI_FITC_TXRED_Cy5-420-450/506-532/578-610/666-724; Dichroic, DAPI_FITC_TXRED_Cy5-415/490/ 572/660) and a CYR71010 filter (Leica : LED, CFP_YFP_RFP_NIR-440/510/575/730; Excitation Range, CFP_YFP_RFP_NIR-422-450/495-517/566-590/710-750; Emission Range, CFP_YFP_RFP_NIR-462-484/527-551/ 602-680/770-850; Dichroic, CFP_YFP_RFP_NIR-459/523/598/763), an LED light source LED8 (Leica microsystems) and DIC to observe the EGFP and mCherry fluorescent signals and hyphal morphology of *A. oryzae* cells. After noise subtraction of the localization data taken using Thunder Imager, analysis was performed using ImageJ Fiji.

### Single molecule fluorescence in situ hybridization (smFISH)

smFISH analysis was performed essentially according to the manufacturer’s instructions (LGC Biosearch Technologies) with some modifications based on the previous study.^54^ Two hundred µL of CD (mal) medium sterilized with a 0.2 µm filter was added to polylysine-coated glass bottom dishes (Matsunami) and inoculated with 1.0 x 10= conidia/10 µl of the *glaA*-36 x MBS and *glaA*-MS2 strains and incubated at 30°C for 24 h. The smFISH probes were used according to the instructions of the manufacturer (LGC Biosearch Technologies). The smFISH probe for 12×*mbs* consisted of mixtures of 18–22 nt from 29 regions in 828 bp, each region was linked to fluorescein amidite (FAM; excitation/emission, 495/520 nm). The smFISH probe for *glaA* consisted of mixtures of 18–22 nt from 48 regions in 1,839 bp, each region was linked to CAL Fluor Red 610 (excitation/emission, 590/610 nm). The information on the 12×*mbs* and *glaA* probe sequences is summarized in Tables S1 and S2.

### Live-cell imaging of *glaA* mRNA by MS2 system

Two hundred µL of CD medium or CD (mal) medium sterilized with 0.2 µm filters was added to polylysine-coated glass bottom dishes and inoculated with 8.0 x 10= conidia/8 µl of *glaA*-MS2 and each co-localization analysis strain and cultured for 20 h or 24 h at 30°C. For microscopy, images were taken with EGFP (FITC, 100%, 100 ms) and mCherry (RFP, 100%, 100 ms). Localization analysis images were processed by the THUNDER imaging system (Leica) with the following settings. For all observations, the inclusion agent in Instant Computational Clearing (ICC) was set to water. In addition, the Feature Scale value of the Advanced Setting in ICC was processed with the following settings: *glaA* mRNA, EGFP=0.4 µm; AoSec61-mCherry, mCherry-AoRab5, AoEdc3-mCherry and AoDcp2-mCherry, mCherry=0.4 µm; mCherry-NLS, mCherry=1.0 µm; AoBre5-mCherry and AoPab1-mCherry, mCherry=1.5 µm.

### Quantitative localization analysis of *glaA* mRNA

*glaA* mRNA counts were measured using the RS-FISH plugin in ImageJ Fiji.^91^ The *glaA* mRNA localization and nuclear localization analysis images were converted from 16 bit to 8 bit and merged, and each hyphal region was cropped. Then, each image was re-segmented and the nuclear localization analysis image was binarized and subtracted from the *glaA* mRNA localization analysis image using the image calculator. The background fluorescence of the *glaA* mRNA localization analysis image, in which nuclear fluorescence was eliminated, was completely subtracted by the Math function and processed with a Gaussian filter. Measurements with RS-FISH were performed with the following settings. In Radial Symmetry, Mode=Interactive, ZYX=1.000, RANSAC was set, Use anisotropy coefficient for Dog was checked, and Spot intensity was set to Linear Interpolation. Then, *glaA* mRNA counts were measured with Adjust difference-of-gaussian values set to sigma=1.5 and Threshold=0.00102.

Correlation coefficients between *glaA* mRNA and ER localization were calculated using the colocalization Finder plug-in. The *glaA* mRNA and ER localization analysis images were converted from 16 bit to 8 bit and merged, and each hyphal region was cropped. Then, the background fluorescence of each image was completely subtracted by the Math function, and the co-localization areas of the *glaA* mRNA and ER localization analysis images were extracted by the image calculator and processed by Gaussian filter. The correlation coefficient was calculated by plotting the co-localization fluorescence of *glaA* mRNA and *glaA* mRNA & ER localization analysis images using the colocalization Finder. As for the ER, regions without nuclei were cropped from the ER localization analysis image. Then, the cytoplasmic ER was extracted by cropping approximately 0.7 µm inward from both sides of the plasma membrane, and the background fluorescence of each image was completely subtracted with the Math function and processed with a Gaussian filter. Total ER and cytoplasmic ER were binarized and the area ratio of total ER to cytoplasmic ER was determined by calculating the area of each image using Analyze Particles.

For Mander’s test of *glaA* mRNA and ER colocalization, *glaA* mRNA and nuclear localization analysis images were converted from 16 bit to 8 bit and merged, and each hyphal region was cropped. Then, each image was re-segmented and the background fluorescence of each image was completely subtracted with the Math function and processed with a Gaussian filter. The *glaA* mRNA and the nuclear localization analysis images were binarized, and the co-localization areas between the *glaA* mRNA and the ER localization analysis images were extracted using the image calculator. Analyze Particles was used to calculate the areas of localization analysis images of *glaA* mRNA, ER and *glaA* mRNA & ER, and the area ratio of localization analysis images of *glaA* mRNA and ER to those of *glaA* mRNA & ER.

### Tracking of *glaA* mRNA dynamics

Observations of *glaA* mRNA were taken with EGFP (FITC, 100%, 20 ms) for 20 sec. Localization analysis of *glaA* mRNA and AoSec61-mCherry (ER) was performed with EGFP (FITC, 100%, 10 ms) and mCherry (TXRED, 100%, 20 ms), respectively, for 30 sec. In similar analyses for the molecular motor deletion strains, EGFP (FITC, 100%, 20 ms) and mCherry (TXRED, 100%, 20 ms) were taken for 20 sec. *glaA* mRNA and mCherry-AoRab5 (EE) were captured with EGFP (FITC, 100%, 20 ms) and mCherry (TXRED, 100%, 20 ms), respectively, for 20 sec. Kinetics of mCherry-AoRab5 (EE) only were captured with mCherry (TXRED, 100%, 20 ms) for 20 sec. Each time-lapse image was processed using the THUNDER imaging system described above. Then, ImageJ Fiji was used and *glaA* mRNA dynamics was analyzed using the TrackMate plug-in. The Dog Detector was used and the threshold was set to 0.3 µm to detect only intracellular-specific fluorescence. Thereafter, Lap Tracker was run with the distance between frames set to 0.5 µm, the gap distance between the three frames set to 0.5 µm and the MAX distance threshold set to 0.8 µm or greater. Co-kinetic analysis of *glaA* mRNA and AoSec61-mCherry (ER) was performed as in the analysis of *glaA* mRNA dynamics. The kymographs for *glaA* mRNA and mCherry-AoRab5 (EE) kinetics were done using the kymograph Builder plugin.

### Inhibitor experiments

To inhibit transcription, Actinomycin D (Act D, stock concentration of 100 mg/ml in DMSO) was used. Sterile 200 µl CD medium with 0.2 µm filter was added to a polylysine-coated glass bottom dish, inoculated with 8.0 x 10= conidia/8 µl of the *glaA*-MS2 strain and incubated at 30°C for 20 h. Then, Act D was added at a final concentration of 100 µg/ml and incubated at 30°C for 1 h. The cells were further incubated for 2 h at 30°C in CD (mal) medium supplemented with Act D at the same final concentration. To restore the transcription of *glaA* mRNA repressed by Act D, the cells were incubated for 2 h at 30°C in CD (mal) medium without Act D. In experiments to confirm transcription inhibition, the PaG strain expressing cytoplasmic EGFP was inoculated at 8.0 x 10= conidia/8 µl and incubated at 30°C for 20 h in CD (glycerol) medium. Then, the cells were incubated for 1 h at 30°C in CD (glycerol) medium supplemented with Act D to the same final concentration. The cells were further incubated in CD (mal) medium supplemented with Act D for 2.5 h at 30°C. The EGFP brightness was line-scanned using the ImageJ Fiji plot profile.

To inhibit microtubule polymerization, nocodazole (Noc, stock concentration of 10 mg/mL in DMSO) was used. Sterile 200 µl CD (mal) medium with 0.2 µm filter was added to a polylysine-coated glass bottom dish, inoculated with 8.0 x 10= conidia/8 µl of the *glaA*-MS2 strain and incubated at 30°C for 24 h. Subsequently, Noc was added at a final concentration of 100 µg/mL and incubated at 30°C for 30 min. As a negative control experiment, observations were also made under conditions in which the same amount of DMSO was added. To confirm the inhibition of microtubule polymerization, the NSlDSER5 strain expressing the EE marker EGFP-AoRab5 was inoculated 8.0 x 10= conidia/8 µl in CD medium and incubated at 30°C for 20 h. Then, Noc was added at the same final concentration and incubated at 30°C for 30 min. The EE kinetics were analyzed using ImageJ Fiji’s kymograph Builder.

To inhibit actin polymerization, latrunculin B (Lat B, stock concentration of 10 mM in DMSO) was used. Sterile 200 µl CD (mal) medium with 0.2 µm filter was added to a polylysine-coated glass bottom dish, inoculated with 8.0 x 10= conidia/8 µl of the *glaA*-MS2 strain and incubated at 30°C for 24 h. Then, Lat B was added to reach a final concentration of 100 µM and incubated at 30°C for 30 min. As a negative control experiment, observations were also made under conditions in which the same amount of DMSO was added. To confirm inhibition of actin polymerization, the PaLaG strain expressing Lifeact-EGFP was inoculated into CD medium at 8.0 x 10= conidia/8 µl and incubated at 30°C for 20 h. Then, Lat B was added at the same final concentration and incubated at 30°C for 30 min. The Lifeact-EGFP fluorescence was line-scanned using ImageJ Fiji’s plot profile. Similarly, the AoAbp1-EGFP expressing strain, abE1, which visualizes an actin-binding protein, was cultured in M medium to confirm inhibition of actin polymerization.

To induce ER stress, dithiothreitol (DTT, stock concentration of 1 M in water) was used. Sterile 200 µl CD (mal) medium with 0.2 µm filter was added to a polylysine-coated glass bottom dish, inoculated with 8.0 x 10= conidia/8 µl of the *glaA*-MS2 strain and incubated at 30°C for 20 h. Then, DTT was added at a final concentration of 10 mM and incubated at 30°C for 2 h. For the measurement of *glaA* mRNA counts, the *glaA*-MS2 strain was inoculated with 8.0 x 10= conidia/8 µl in CD (mal) medium and incubated at 30°C for 20 h. Then, DTT was added at the same final concentration and incubated at 30°C for 1 h. To recover from ER stress, the cells were incubated in CD (mal) medium without DTT at 30°C for 2 h. As a negative control experiment, observations were also made under conditions in which the same amount of water was added.

### Quantification and statistical analysis

All experiments were conducted at least twice. Quantitative results are shown as mean ± SD. Statistical analysis was performed by Student’s *t*-test or Tukey-Kramer test with the quantitative data.

## SUPPLEMENTAL INFORMATION

Supplemental information includes 6 Supplemental figures, 2 Supplemental tables and 5 Supplemental movies.

## Notes

### Competing Interest Statement

The authors have declared no competing interest.

