## Supplemental information for "Polarity-dependent expression and localization of secretory glucoamylase mRNA in filamentous fungal cells"

Word count: Summary, 150; Main, 6728.

Number of figures: Main, 7; Supplemental, 6.

Number of tables: Supplemental, 2.

Number of movies: Supplemental, 5.

#### SUPPLEMENTAL FIGURE LEGENDS

**Figure S1. Effects on growth, expression and activity of the *glaA*-MS2 strain**

(A) Control (NSIDS1) and *glaA*-MS2 strains were grown on Czapek-Dox (CD) agar plates containing potato starch (PS) as a sole carbon source and DPY agar plates at 30°C for 3 days.

(B) Diameters of the colony were determined at day 3 for each culture condition. Error bars indicate the standard deviation of the mean (n = 3).

(C) Relative expression of *glaA* mRNA. Control (NSIDS1) and *glaA*-MS2 strains were grown in CD (mal) medium at 30°C for 24 h, and then the relative expression of *glaA* was determined. Error bars indicate the standard deviation of the mean (n = 3).

(D and E) Control (NSIDS1) and *glaA*-MS2 strains were grown in DPY liquid medium at 30°C for 3 days, and then dry mycelium weight and glucoamylase activity were determined. Error bars indicate the standard deviation of the mean (n = 3).

**Figure S2. Validation tests of transcription inhibition by Actinomycin D**

(A) Comparison of EGFP intensity in the PaG strain expressing cytoplasmic EGFP. The strain was grown in CD medium containing glycerol as a sole carbon source at 30°C for 20 h. After that, cultures were further incubated in CD medium containing glycerol in the presence of DMSO or ActD at 30°C for 1 h.

(B) After culturing in CD medium containing glycerol overnight, cells were incubated in CD medium containing maltose to induce the *glaA* expression in the presence of DMSO or ActD at 30°C for 2.5 h.

(C) Line scan analysis of the EGFP intensity on the blue lines in (B) for each condition was performed by plot profile of ImageJ Fiji.

**Figure S3. Localization of *glaA* mRNA and ER**

(A) Localization of *glaA* mRNAs (orange) and nuclei (blue) by smFISH in the NSIDN1 strain. *glaA* mRNA was labeled by the *glaA* specific probe. The white and yellow arrows indicate the hyphal tip and conidium, respectively. Scale bars, 10 µm.

(B) Number of nuclei in each hyphal area was determined in the *glaA*-MS2-N strain.

Error bars indicate the standard deviation of the mean (n = 10). Significant difference at P < 0.05 between a and b (Tukey–Kramer test).

(C) Localization of *glaA* mRNAs (green) and nuclei (magenta) in a branch point (red arrows) of the basal region. Scale bars, 10 μm.

(D) Localization of *glaA* mRNAs (green) and nuclei (magenta) in a wide area. The culture was shifted from CD to CD (mal) media to induce *glaA* mRNA expression. White, blue and yellow arrows indicate the tip, septum and conidium, respectively. Scale bars, 50 μm.

(E) A phylogenetic tree of AoSec61 was made by using BLAST of GenomeNet (https://www.genome.jp/tools/blast/).

(F) Localization of *glaA* mRNAs (green) and ER (AoSec61-mCherry, magenta). White, blue and yellow arrows indicate the tip, septum and conidium, respectively. Scale bars, 10 μm.

**Figure S4. Validation tests of polymerization inhibition of actin and microtubule**

(A and B) Localization of AoAbp1-EGFP and Lifeact-EGFP in the preculture (pre) or in the presence of DMSO or latrunculin B. White arrows show the hyphal tip. Scale bars, 10 μm.

(C) Line scan analysis of the Lifeact-EGFP intensity on the blue lines in (B) for each condition was performed by plot profile of ImageJ Fiji.

(D) Kymographs of EGFP-AoRab5 dynamics in the preculture (pre) or in the presence of DMSO or nocodazole were generated in the regions of the white double arrows by using Kymograph Builder of ImageJ Fiji. White arrows show the hyphal tip. Scale bars, 10 μm.

**Figure S5. Effects on EE dynamics and growth by kinesin motor disruptants**

(A and B) Phylogenetic trees of AoKin1 and AoKin3 were made by using BLAST of GenomeNet (https://www.genome.jp/tools/blast/).

(C and D) Kymographs of EE (mCherry-AoRab5) dynamics in the Δ*Aokin1* and Δ*Aokin3* strains were generated in the regions of the white double arrows by using

Kymograph Builder of ImageJ Fiji. White arrows show the hyphal tip. Scale bars, 10 μm.

(E) Growth assay of the *glaA*-MS2, Δ*Aokin1*, Δ*Aokin3* and Δ*Aokin1*Δ*Aokin3* strains. The schematic shows the spotted position of each strain. These strains were grown on CDm agar plates containing either glucose, maltose or potato starch as a sole carbon source and a DPY agar plate at 30°C for 3 days.

(F to I) Diameters of the colony grown on CDm (F), CDm (mal), CDm (PS) and DPY were determined at day 3 for each culture condition. Error bars indicate the standard deviation of the mean (n = 3). *** Statistically significant difference at $P < 0.001$.

**Figure S6. Colocalization of *glaA* mRNA, SG and PB**

(A and B) Phylogenetic trees of AoBre5 and AoEdc3 were made by using BLAST of GenomeNet (https://www.genome.jp/tools/blast/).

(C to E) Colocalization of *glaA* mRNAs (green), SG (magenta, AoPab1-mCherry; green, AoPab1-EGFP) and PB (magenta, AoDcp2-mCherry) at normal, 45°C for 10 min and 45°C for 30 min. White arrows show the hyphal tip and white arrowheads show colocalization sites. Scale bars, 10 μm.

### Figure S1. Morita et al.

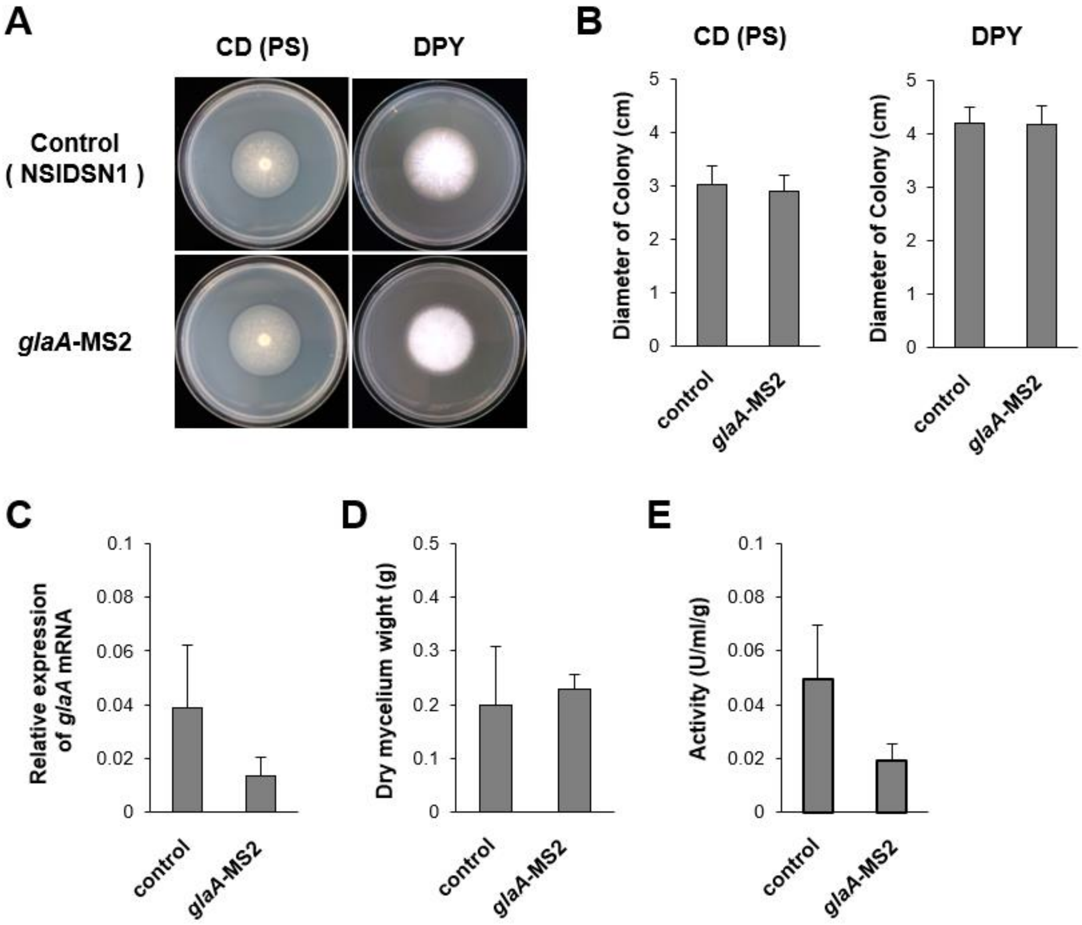

### Figure S2. Morita et al.

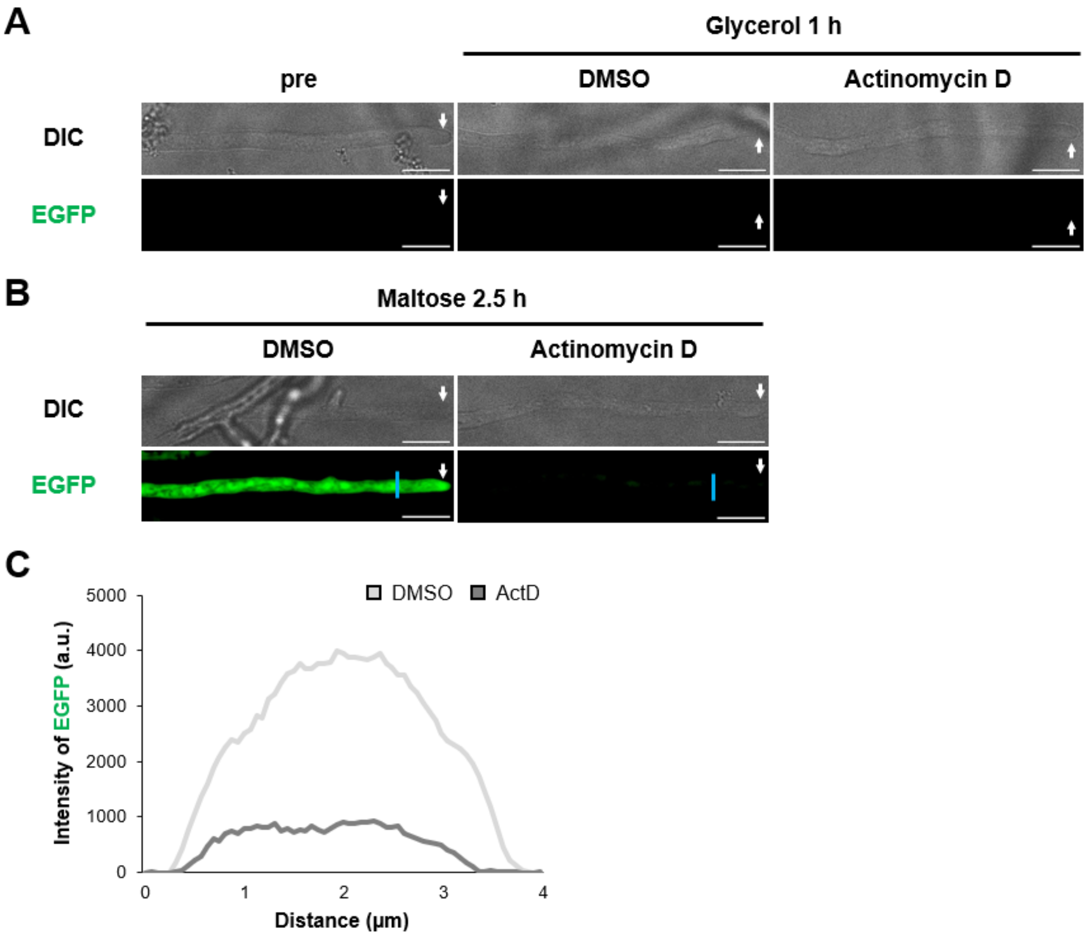


97

**Figure S3. Morita et al.**

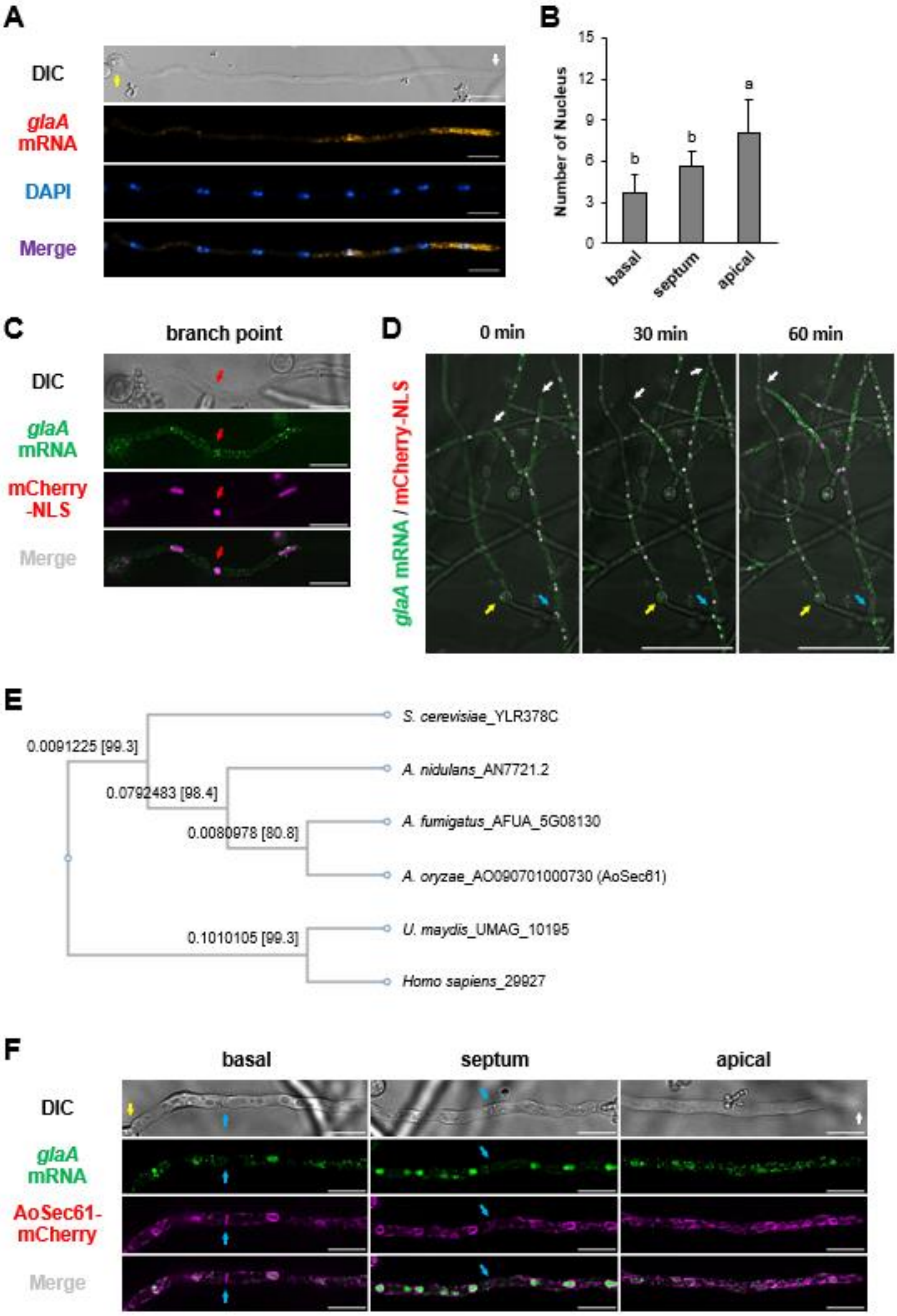


98

### Figure S4. Morita et al.

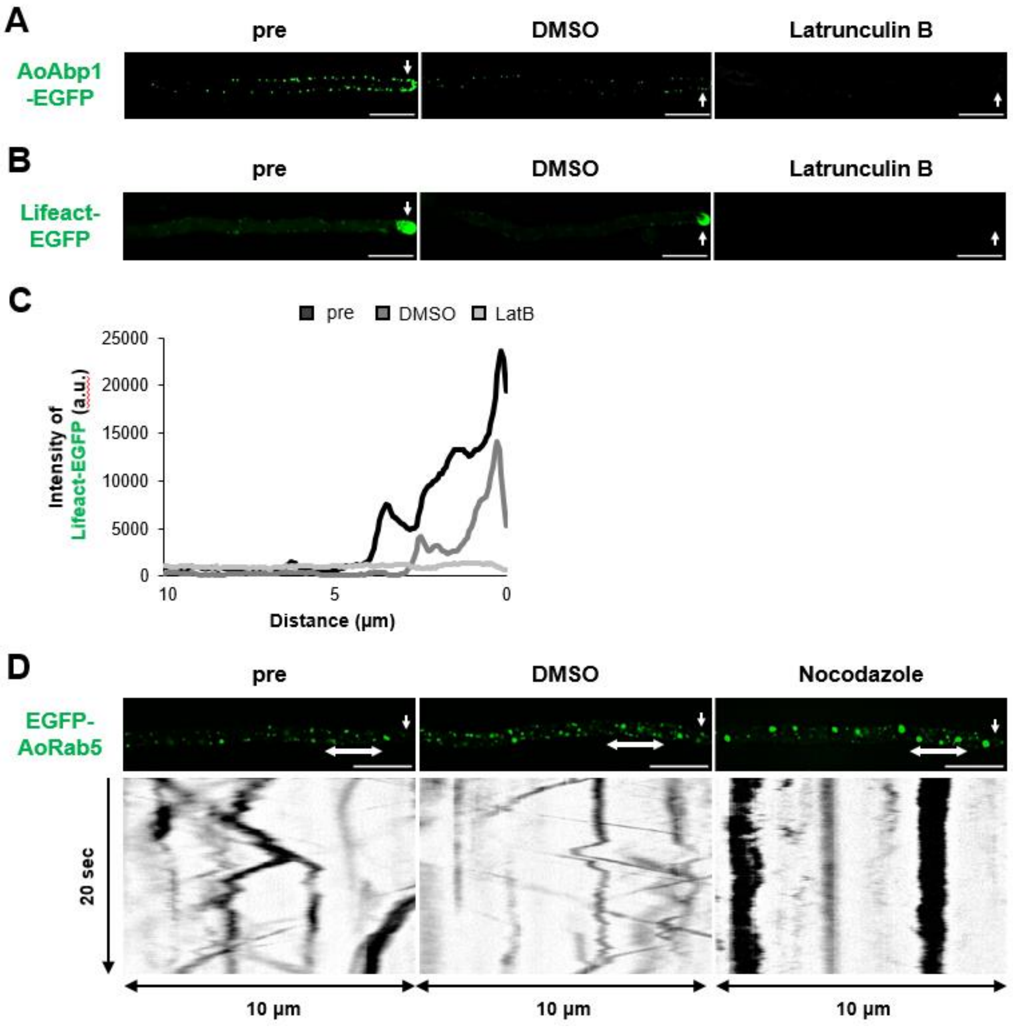

#### Figure S5. Morita et al.

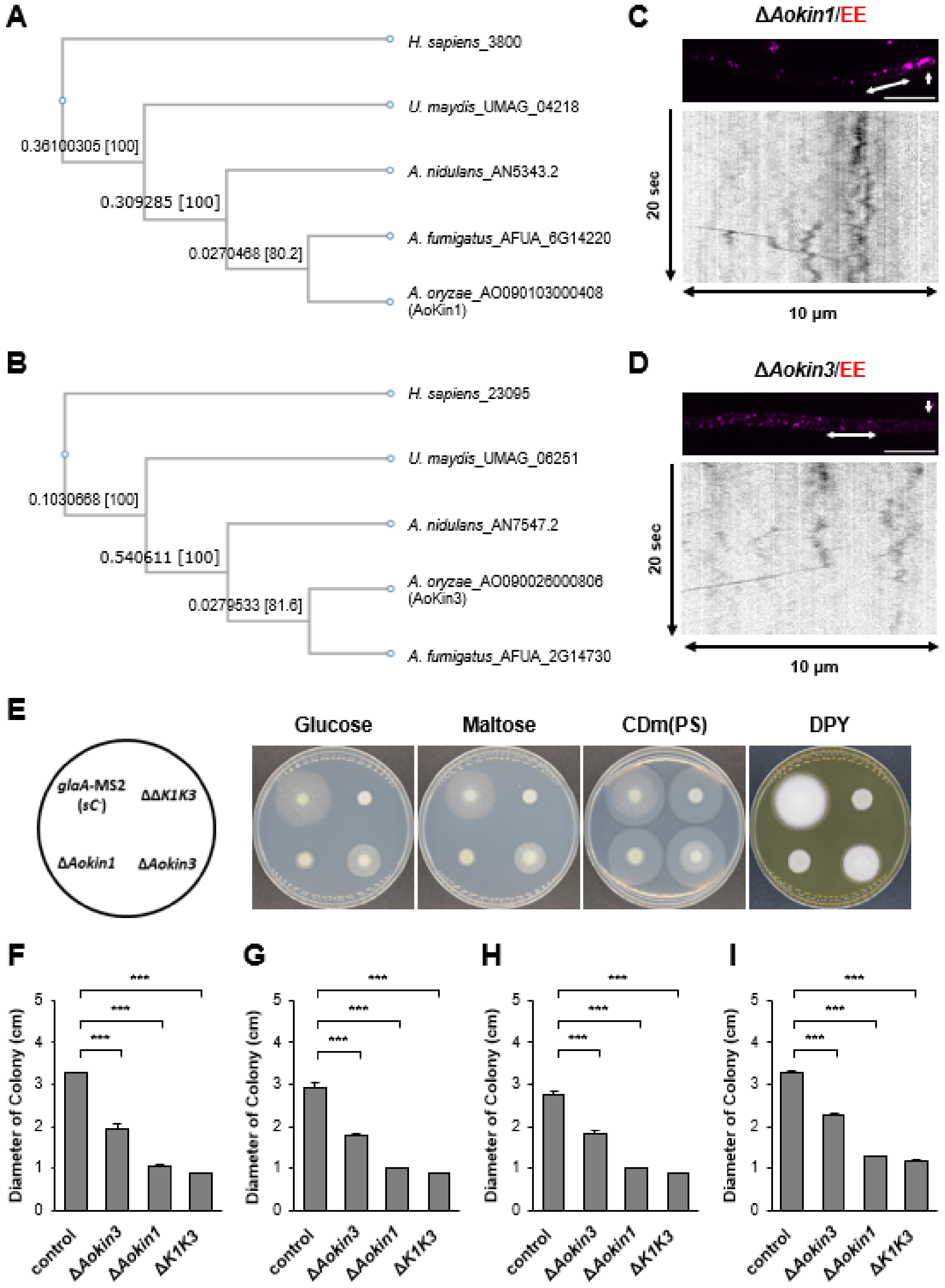

#### Figure S6. Morita et al.

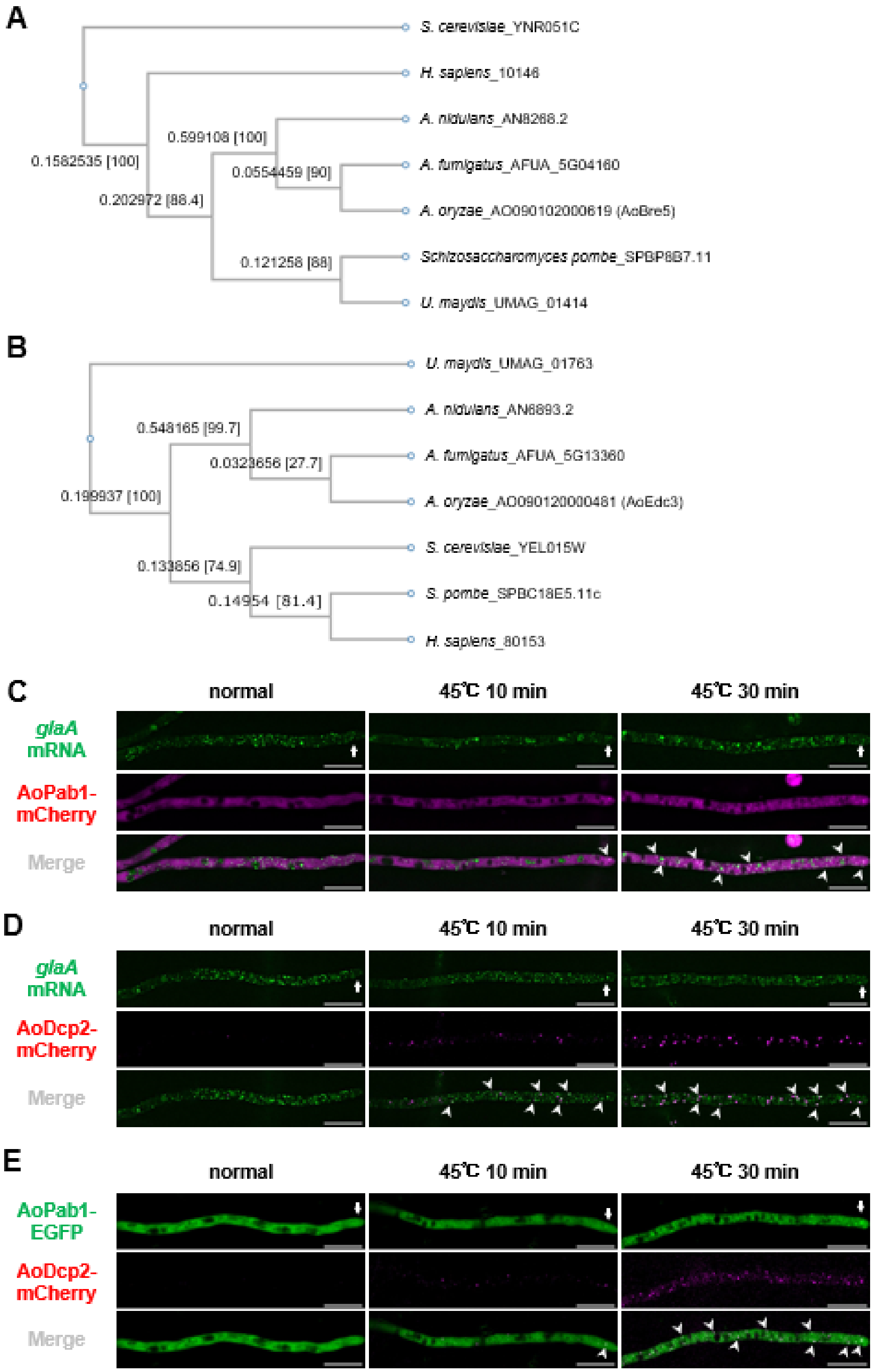

#### SUPPLEMENTAL TABLE

##### Table S1. 12×*mbs* probe sequence used for smFISH

```
ggtattctgccgatgtacgagaagacgattacgcttctcgactaccttcatcatacatgg
cc  aagacggctacatgctcttc  ctaatgcgaagagctgatgg   tagtatgtacc
    Probe # 15            Probe # 16             Probe # 17

tgtgcagatggccgccaagtttttgccaatggaggaataccccattctctgtcaaccaa
acacgtcta            aaaacggttacctccttatg     agagacagttggtt
                     Probe # 18               Probe # 19

cccatgcaaagtttacactctgctatggcagcactgtcgcagaggaataccctgcgagcc
gggtac   tcaaatgtgagacgataccg                  cttatgggacgctcgg
         Probe # 20                            Probe # 21

aaaacggcccccggtgcgtgtattgcatctgccttgcgagcattcacagggacgaatacg
tttt                acataacgtagacggaacgc  gtaagtgtccctgcttatgc
                    Probe # 22            Probe # 23

ccctgccgttgcattacttcaatatgggtgctctgtcgtcgtcatcaggaccatttgcgc
              atgaagttatacccacgaga  gcagcagtagtcctggtaaa   g
              Probe # 24            Probe # 25             P

aggactaccgcgcatatatcatcagcactcgtgcggatacttctgggattcctattgtta
tcctgatggcgcgtatata   gtcgtgagcacgcctatgaa  ccctaaggataacaat
robe # 26             Probe # 27            Probe # 28

cgcgagctcaggaataccgagctctggcgacagagaccctcacacgga
gcgc  gagtccttatggctcgagac
      Probe # 29
```

```
cagagccccctggcaatcgcggggagacgcaggactaccgcgtcttcactctcgcttgcg
                                        gcagaagtgagagcgaacgc
                                        Probe # 1

cggtatacgcagggagcaacgtcacccagggcgacacagaggaataccctgtgtggcccc
                                  gtgtctccttatgggacaca
                                  Probe # 2

acggtgctacaaagaacattccgtattgctcaccagtgtcacatgtggaggactacccca
 gccacgatgtttcttgtaag    aacgagtggtcacagtgtac
 Probe # 3               Probe # 4

caagcgagtcaggaaaccttcggggcatcgcaccattatcccgaacaatcgacacgcagg
gttcgctcagtcctttggaa    cgtagcgtggtaatagggct  ttagctgtgcgtcc
Probe # 5               Probe # 6             Probe # 7

attaccgcgtgggacactctgttccccgtcaaaattggaccataccggagtcggcttacg
taatgg       tgtgagacaaggggcagttt                  ctcagccgaatgc
             Probe # 8                             Probe # 9

tcatgcaggattaccgcatgcatggtgcagaatatcggcatgtctactcgtacagtcaaa
agtacgt  taatggcgtacgtaccacgt  tatagccgtacagatgagca  tcagttt
         Probe # 10            Probe # 11            Probe #

tctactcgtgtggtcaggactaccgaccacttctattctattcatcttttcggttgtgca
agatgagcacacc   cctgatggctggtgaagata      gtagaaaagccaacacgt
 12             Probe # 13                Probe # 14
```

##### Table S2. *glaA* probe sequence used for smFISH

```
[illegible]
Probe # 1   Probe # 2   Pro

[illegible]
be # 3   Probe #

[illegible]
4   Probe # 5

[illegible]
Probe # 6   Probe # 7

[illegible]
Probe # 8   Probe #

[illegible]
9   Probe # 10   Probe

[illegible]
# 11   Probe # 12

[illegible]
Probe

[illegible]
# 13   Probe # 14

[illegible]
Probe # 15   Probe # 16   Probe # 1

[illegible]
7   Probe # 18   Probe #
```

```
[illegible]
19   Probe # 20   Probe # 21

[illegible]
Probe # 22   Probe # 23

[illegible]
Probe # 24   Probe # 25

[illegible]
Probe # 26

[illegible]
Probe # 27   Probe # 28

[illegible]
Probe # 29   Probe # 30   P

[illegible]
robe # 31   Probe # 32   Pr

[illegible]
obe # 33   Probe # 34   Probe

[illegible]
# 35   Probe # 36   Probe #

[illegible]
37   Probe # 38

[illegible]
Probe # 39

[illegible]
Probe # 40
```

```
[illegible]
Probe # 41

[illegible]
Probe # 42

[illegible]
Probe # 4

[illegible]
3   Probe # 44

[illegible]
Probe # 45

[illegible]
Probe # 46   Probe # 47

[illegible]
Probe # 48

[illegible]
```

#### SUPPLEMENTAL MOVIE LEGENDS

**Movie S1. Co-dynamics of *glaA* mRNA and ER in the *glaA*-MS2 strain**

Co-dynamics of *glaA* mRNAs (green) and ER (magenta, AoSec61-mCherry). The strain was grown in CD (mal) medium at 30°C for 24 h. Time-lapse images are shown for 30 sec (7.23 frame/sec). Scale bar, 10 μm.

**Movie S2. Co-dynamics of *glaA* mRNA and EE in the *glaA*-MS2 strain**

Co-dynamics of *glaA* mRNAs (green) and EE (magenta, mCherry-AoRAb5). The strain was grown in CD (mal) medium at 30°C for 24 h. Time-lapse images are shown for 20 sec (8.45 frame/sec). Scale bar, 10 μm.

**Movie S3. Co-dynamics of *glaA* mRNA and ER in the Δ*Aokin3* strain**

Co-dynamics of *glaA* mRNAs (green) and ER (magenta, AoSec61-mCherry). The strain was grown in CD (mal) medium at 30°C for 24 h. Time-lapse images are shown for 20 sec (6.75 frame/sec). Scale bar, 10 μm.

**Movie S4. Co-dynamics of *glaA* mRNA and ER in the Δ*Aokin1* strain**

Co-dynamics of *glaA* mRNAs (green) and ER (magenta, AoSec61-mCherry). The strain was grown in CD (mal) medium at 30°C for 24 h. Time-lapse images are shown for 20 sec (6.75 frame/sec). Scale bar, 10 μm.

**Movie S5. Co-dynamics of *glaA* mRNA and ER in the Δ*Aokin1*Δ*Aokin3* strain**

Co-dynamics of *glaA* mRNAs (green) and ER (magenta, AoSec61-mCherry). The strain was grown in CD (mal) medium at 30°C for 24 h. Time-lapse images are shown for 20 sec (6.75 frame/sec). Scale bar, 10 μm.
