## Supplementary material for "Polarity-dependent expression and localization of secretory glucoamylase mRNA in filamentous fungal cells": Resource table

| REAGENT or RESOURCE | SOURCE | IDENTIFIER |
|---|---|---|
| Bacterial and virus strains | | |
| *Escherichia coli* DH5α Competent Cells | TaKaRa | Cat#9057 |
| Chemicals, peptides, and recombinant proteins | | |
| DMSO | FUJIFILM Wako | Cat#049-07213 |
| Actinomycin D (LatB) | FUJIFILM Wako | Cat#010-21263 |
| Nocodazole (Noc) | Sigma | Cat#31430-18-9 |
| Latrunculin B (LatB) | Tocris Bioscience | Cat#3974 |
| Dithiothreitol (DTT) | FUJIFILM Wako | Cat#042-29222 |
| Experimental models: Organisms/strains | | |
| *A. oryzae* RIB40 (WT) | N/A | N/A |
| NSlDSN1 (*niaD*$^-$ *niaD sC*$^-$ *AosC adeA*$^-$ Δ*argB*::*adeA*$^-$ Δ*ligD*::*argB* Δ*pyrG*::*adeA pyrG*) | Togo et al.[46] | N/A |
| NSlDN1 (*niaD*$^-$ *niaD sC*$^-$ *adeA*$^-$ Δ*argB*::*adeA*$^-$ Δ*ligD*::*argB* Δ*pyrG*::*adeA pyrG*) | Morita et al.[92] | N/A |
| abE1 (*niaD*$^-$ *sC*$^-$ *Aoabp1-egfp*::*AosC adeA*$^-$ *adeA* Δ*argB* Δ*ku70*::*argB*) | This study | N/A |
| NSlDSER5 (*niaD*$^-$ P*amyB-egfp-Aorab5 niaD sC*$^-$ *AosC adeA*$^-$ Δ*argB::adeA*$^-$ Δ*ligD::argB* Δ*pyrG::adeA pyrG*) | Togo et al.[46] | N/A |
| PaG (*niaD*$^-$ P*amyB-egfp*::*niaD sC*$^-$ *AosC adeA*$^-$ Δ*argB*::*adeA*$^-$ Δ*ligD*::*argB* Δ*pyrG*::*adeA pyrG*) | Morita et al.[92] | N/A |
| PaLaG (*niaD*$^-$ P*amyB-Lifeact-egfp*::*niaD sC*$^-$ *AosC adeA*$^-$ Δ*argB*::*adeA*$^-$ Δ*ligD*::*argB* Δ*pyrG*::*adeA pyrG*) | Morita et al.[92] | N/A |
| PpP1GD2mCh (*niaD*$^-$ P*pgkA-Aopab1-egfp*::*niaD sC*$^-$ P*pgkA-Aodcp2-egfp*::*sC adeA*$^-$ Δ*argB*::*adeA*$^-$ Δ*ligD*::*argB* Δ*pyrG*::*adeA pyrG*) | This study | N/A |
| *glaA*-36×MBS (*niaD*$^-$ *sC*$^-$ *adeA*$^-$ Δ*argB*::*adeA*$^-$ Δ*ligD*::*argB* Δ*pyrG*::*adeA*) | This study | N/A |
| *glaA*-MS2 (*niaD*$^-$ P*pgkA-nls-mcp-2×egfp*::*niaD sC*$^-$ *AosC adeA*$^-$ Δ*argB*::*adeA*$^-$ Δ*ligD*::*argB* Δ*pyrG*::*adeA pyrG*) | This study | N/A |
| *glaA*-MS2-N (*niaD*$^-$ P*pgkA-nls-mcp-2×egfp*::*niaD sC*$^-$ P*Aotps1-mcherry-nls*::*AosC adeA*$^-$ Δ*argB*::*adeA*$^-$ Δ*ligD*::*argB* Δ*pyrG*::*adeA pyrG*) | This study | N/A |
| *glaA*-MS2-S61 (*niaD*$^-$ P*pgkA-nls-mcp-2×egfp*::*niaD sC*$^-$ P*pgkA-Aosec61-mcherry*::*AosC adeA*$^-$ Δ*argB*::*adeA*$^-$ Δ*ligD*::*argB* Δ*pyrG*::*adeA pyrG*) | This study | N/A |
| *glaA*-MS2-R5 (*niaD*$^-$ P*pgkA-nls-mcp-2×egfp*::*niaD sC*$^-$ P*pgkA-mcherry-Aorab5*::*AosC adeA*$^-$ Δ*argB*::*adeA*$^-$ Δ*ligD*::*argB* Δ*pyrG*::*adeA pyrG*) | This study | N/A |
| *glaA*-MS2-B5 (*niaD*$^-$ P*pgkA-nls-mcp-2×egfp*::*niaD sC*$^-$ P*pgkA-Aobre5-mcherry*::*AosC adeA*$^-$ Δ*argB*::*adeA*$^-$ Δ*ligD*::*argB* Δ*pyrG*::*adeA pyrG*) | This study | N/A |
| *glaA*-MS2-E3 (*niaD*$^-$ P*pgkA-nls-mcp-2×egfp*::*niaD sC*$^-$ P*pgkA-Aoedc3-mcherry*::*AosC adeA*$^-$ Δ*argB*::*adeA*$^-$ Δ*ligD*::*argB* Δ*pyrG*::*adeA pyrG*) | This study | N/A |
| *glaA*-MS2-P1 (*niaD*$^-$ P*pgkA-nls-mcp-2×egfp*::*niaD sC*$^-$ P*pgkA-Aopab1-mcherry*::*AosC adeA*$^-$ Δ*argB*::*adeA*$^-$ Δ*ligD*::*argB* Δ*pyrG*::*adeA pyrG*) | This study | N/A |

| | | |
|---|---|---|
| *glaA*-MS2-D2 (*niaD*⁻ P*pgkA-nls-mcp-2×egfp*::*niaD sC*⁻ P*pgkA-Aodcp2-mcherry*::*AosC adeA*⁻ Δ*argB*::*adeA*⁻ Δ*ligD*::*argB* Δ*pyrG*::*adeA pyrG*) | This study | N/A |
| *glaA*-MS2-ΔH1 (*niaD*⁻ P*pgkA-nls-mcp-2×egfp*::*niaD sC*⁻ *adeA*⁻ Δ*argB*::*adeA*⁻ Δ*ligD*::*argB* Δ*pyrG*::*adeA* Δ*Aohok1*::*pyrG*) | This study | N/A |
| *glaA*-MS2-ΔH1N (*niaD*⁻ P*pgkA-nls-mcp-2×egfp*::*niaD sC*⁻ *adeA*⁻ P*Aotps1-mcherry-nls*::*AosC* Δ*argB*::*adeA*⁻ Δ*ligD*::*argB* Δ*pyrG*::*adeA* Δ*Aohok1*::*pyrG*) | This study | N/A |
| *glaA*-MS2-ΔH1S61 (*niaD*⁻ P*pgkA-nls-mcp-2×egfp*::*niaD sC*⁻ *adeA*⁻ P*pgkA-Aosec61-mcherry*::*AosC* Δ*argB*::*adeA*⁻ Δ*ligD*::*argB* Δ*pyrG*::*adeA* Δ*Aohok1*::*pyrG*) | This study | N/A |
| *glaA*-MS2-ΔK1 (*niaD*⁻ P*pgkA-nls-mcp-2×egfp*::*niaD sC*⁻ *adeA*⁻ Δ*argB*::*adeA*⁻ Δ*ligD*::*argB* Δ*pyrG*::*adeA* Δ*Aokin1*::*pyrG*) | This study | N/A |
| *glaA*-MS2-ΔK1N (*niaD*⁻ P*pgkA-nls-mcp-2×egfp*::*niaD sC*⁻ P*Aotps1-mcherry-nls*::*AosC adeA*⁻ Δ*argB*::*adeA*⁻ Δ*ligD*::*argB* Δ*pyrG*::*adeA* Δ*Aokin1*::*pyrG*) | This study | N/A |
| *glaA*-MS2-ΔK1S61 (*niaD*⁻ P*pgkA-nls-mcp-2×egfp*::*niaD sC*⁻ P*pgkA-Aosec61-mcherry*::*AosC adeA*⁻ Δ*argB*::*adeA*⁻ Δ*ligD*::*argB* Δ*pyrG*::*adeA* Δ*Aokin1*::*pyrG*) | This study | N/A |
| *glaA*-MS2-ΔK1R5 (*niaD*⁻ P*pgkA-nls-mcp-2×egfp*::*niaD sC*⁻ P*pgkA-mcherry-Aorab5*::*AosC adeA*⁻ Δ*argB*::*adeA*⁻ Δ*ligD*::*argB* Δ*pyrG*::*adeA* Δ*Aokin1*::*pyrG*) | This study | N/A |
| *glaA*-MS2-ΔK3 (*niaD*⁻ P*pgkA-nls-mcp-2×egfp*::*niaD sC*⁻ *adeA*⁻ Δ*argB*::*adeA*⁻ Δ*ligD*::*argB* Δ*pyrG*::*adeA* Δ*Aokin3*::*pyrG*) | This study | N/A |
| *glaA*-MS2-ΔK3N (*niaD*⁻ P*pgkA-nls-mcp-2×egfp*::*niaD sC*⁻ P*Aotps1-mcherry-nls*::*AosC adeA*⁻ Δ*argB*::*adeA*⁻ Δ*ligD*::*argB* Δ*pyrG*::*adeA* Δ*Aokin3*::*pyrG*) | This study | N/A |
| *glaA*-MS2-ΔK3S61 (*niaD*⁻ P*pgkA-nls-mcp-2×egfp*::*niaD sC*⁻ P*pgkA-Aosec61-mcherry*::*AosC adeA*⁻ Δ*argB*::*adeA*⁻ Δ*ligD*::*argB* Δ*pyrG*::*adeA* Δ*Aokin3*::*pyrG*) | This study | N/A |
| *glaA*-MS2-ΔK3R5 (*niaD*⁻ P*pgkA-nls-mcp-2×egfp*::*niaD sC*⁻ P*pgkA-mcherry-Aorab5*::*AosC adeA*⁻ Δ*argB*::*adeA*⁻ Δ*ligD*::*argB* Δ*pyrG*::*adeA* Δ*Aokin3*::*pyrG*) | This study | N/A |
| *glaA*-MS2-ΔΔK1K3 (*niaD*⁻ P*pgkA-nls-mcp-2×egfp*::*niaD sC*⁻ *adeA*⁻ Δ*argB*::*adeA*⁻ Δ*ligD*::*argB* Δ*pyrG*::*adeA* Δ*Aokin1*::*ptrA* Δ*Aokin3*::*pyrG*) | This study | N/A |
| *glaA*-MS2-ΔΔK1K3N (*niaD*⁻ P*pgkA-nls-mcp-2×egfp*::*niaD sC*⁻ P*Aotps1-mcherry-nls*::*AosC adeA*⁻ Δ*argB*::*adeA*⁻ Δ*ligD*::*argB* Δ*pyrG*::*adeA* Δ*Aokin1*::*ptrA* Δ*Aokin3*::*pyrG*) | This study | N/A |
| *glaA*-MS2-ΔΔK1K3S61 (*niaD*⁻ P*pgkA-nls-mcp-2×egfp*::*niaD sC*⁻ P*pgkA-Aosec61-mcherry*::*AosC adeA*⁻ Δ*argB*::*adeA*⁻ Δ*ligD*::*argB* Δ*pyrG*::*adeA* Δ*Aokin1*::*AnptrA* Δ*Aokin3*::*pyrG*) | This study | N/A |
| Oligonucleotides | | |
| *gpdA* -Fw-RT (CGTCGAGTCCACTGGTGTCTT) | Morita et al.[92] | N/A |
| *gpdA*-Rv-RT (TTGTTGACACCCATAACGAACATGG) | Morita et al.[92] | N/A |

| | | |
|---|---|---|
| *glaA*-Fw-RT (CGGTTCCTCGTGCCCCTATT) | Togo et al.[46] | N/A |
| *glaA*-Rv-RT (GATGTCTTTGCCCGATCGCC) | Togo et al.[46] | N/A |
| Recombinant DNA | | |
| pgNotI-AosC-NotI (NotI-*AosC*-NotI) | This study | N/A |
| pgSmaI-pyrG-ptrA (SmaI-*pyrG*-*ptrA*) | This study | N/A |
| pET264 (pUC 24xMS2V6 Loxp KANr Loxp) | Tutucci et al.[1] | Addgene #104393 |
| pANPXP-Cre (*lox-AnptrA-Cre-lox*) | Zhang et al.[87] | N/A |
| 36×MBS cassette (NotI-36×*mbs-lox-AnptrA-Cre-lox*-NotI) | This study | N/A |
| glaA-MBS-AnptrA cassette (NotI-*glaA* ORF-*lox-ptrA-Cre-lox-glaA* 3'UTR-NotI) | This study | N/A |
| glaA-MBS cassette (NotI-*glaA* ORF-*lox-pyrG-Cre-lox-glaA* 3'UTR-NotI) | This study | N/A |
| pgPpSmaIG (P*pgkA*-SmaI-*egfp*-T*amyB*-*niaD*) | This study | N/A |
| pgPpSmaImC (P*pgkA*-SmaI-*mcherry*-T*amyB*-*AosC*) | This study | N/A |
| pgPpMG (P*pgkA*-*mcp*-*egfp*-T*amyB*-*niaD*) | This study | N/A |
| pgPpM2G (P*pgkA*-*mcp*-2×*egfp*-T*amyB*-*niaD*) | This study | N/A |
| pgPcNM2G (P*Aocyc1*-*nls*-*mcp*-2×*egfp*-T*amyB*-*niaD*) | This study | N/A |
| pgPpNM2G (P*pgkA*-*nls*-*mcp*-2×*egfp*-T*amyB*-*niaD*) | This study | N/A |
| pgPtmCN (P*Aotps1*-*mcherry*-*nls*-T*amyB*-*AosC*) | This study | N/A |
| pgPpS61mCN (P*pgkA*-*Aosec61*-*mcherry*-T*amyB*-*niaD*) | This study | N/A |
| pgPpS61mCS (P*pgkA*-*Aosec61*-*mcherry*-T*amyB*-*AosC*) | This study | N/A |
| pgPpmCR5 (P*pgkA*-*mcherry*-*Aorab5*-T*amyB*-*AosC*) | This study | N/A |
| pgPpB5mC (P*pgkA*-*Aobre5*-*mcherry*-T*amyB*-*AosC*) | This study | N/A |
| pgPpE3mC (P*pgkA*-*Aoedc3*-*mcherry*-T*amyB*-*AosC*) | This study | N/A |
| pgPpP1G (P*pgkA*-*Aopab1*-*egfp*-T*amyB*-*niaD*) | This study | N/A |
| pgPpP1mC (P*pgkA*-*Aopab1*-*mcherry*-T*amyB*-*AosC*) | This study | N/A |
| pgPpD2mC (P*pgkA*-*Aodcp2*-*mcherry*-T*amyB*-*AosC*) | This study | N/A |
| pgAbp1-EGFP (SpeI-5'*abp1*-T*amyB*-*AosC*-3'*abp1*-SpeI) | This study | N/A |
| ΔAohok1-pyrG cassette (NotI-*Aohok1* 5'UTR-*pyrG*-T*amyB*- *Aohok1* 3'UTR-NotI) | This study | N/A |
| ΔAokin1-pyrG cassette (NotI-*Aokin1* 5'UTR-*pyrG*-T*amyB*- *Aokin1* 3'UTR-NotI) | This study | N/A |
| ΔAokin1-ptrA cassette (NotI-*Aokin1* 5'UTR-*AnptrA*-T*amyB*- *Aokin1* 3'UTR-NotI) | This study | N/A |
| ΔAokin3-pyrG cassette (NotI-*Aokin3* 5'UTR-*pyrG*-T*amyB*- *Aokin3* 3'UTR-NotI) | This study | N/A |
| Software and algorithms | | |
| SHUNDER imaging system | N/A | Leica Microsystems |
| LASX | N/A | Leica Microsystems |
| ImageJ Fiji | Schindelin et al.[93] | https://imagej.net/Fiji |
| Other | | |
| MCP (optimized codon sequence for *A. oryzae*) | This study | Thermo Fisher |
